# *TP53* mutations promote immunogenic activity in breast cancer

**DOI:** 10.1101/354779

**Authors:** Zhixian Liu, Zehang Jiang, Yingsheng Gao, Lirui Wang, Cai Chen, Xiaosheng Wang

## Abstract

**Background:** Although immunotherapy has recently achieved clinical successes in a variety of cancers, thus far there is no any immunotherapeutic strategy for breast cancer (BC). Thus, it is important to discover biomarkers for identifying the BC patients responsive to immunotherapy. *TP53* mutations were often associated with worse clinical outcome in BC, of which the triple-negative BC (TNBC) has a high *TP53* mutation rate (approximately 80%). TNBC is high-risk due to its high invasiveness, and lack of targeted therapy. To explore a potentially promising therapeutic option for the *TP53*-mutated BC subtype, we studied the associations between *TP53* mutations and immunogenic activity in BC.

**Methods:** We compared enrichment levels of 26 immune gene-sets that indicated activities of diverse immune cells, functions, and pathways between *TP53*-mutated and *TP53*-wildtype BCs based on two large-scale BC multi-omics data. Moreover, we explored the molecular cues that were associated with the differences in immunogenic activity between *TP53*-mutated and *TP53*-wildtype BCs. Furthermore, we performed experimental validation of the findings from bioinformatics analysis.

**Results:** We found that almost all analyzed immune gene-sets had significantly higher enrichment levels in *TP53*-mutated BCs compared to *TP53*-wildtype BCs. Moreover, our experiments confirmed that mutant p53 could increase BC immunogenicity. Furthermore, our computational and experimental results showed that *TP53* mutations could promote BC immunogenicity *via* regulation of the p53-mediated pathways including cell cycle, apoptosis, Wnt, Jak-STAT, NOD-like receptor, and glycolysis. Interestingly, we found that elevated immune activities were likely to be associated with better survival prognosis in *TP53*-mutated BCs, but not necessarily in *TP53*-wildtype BCs.

**Conclusions:** *TP53* mutations promote immunogenic activity in breast cancer. This finding demonstrates a different effect of p53 dysfunction on tumor immunogenicity from that of previous studies, suggesting that the *TP53* mutation status could be a useful biomarker for stratifying BC patients responsive to immunotherapy.

## Background

The tumor suppressor p53 plays an important role in cell-cycle regulation, apoptosis, DNA repair, cellular senescence, and autophagy [1]. Accordingly, *TP53* mutations and dysfunction are importantly involved in carcinogenesis due to the disturbance of these biologic processes it functions in. In fact, *TP53* mutations occur in more than half of all human cancer cases [2], and are independent markers of poor prognosis in a variety of cancers [3]. P53 also plays an important role in immune regulation, e.g., the control of immune responses to infection, autoimmunity and cancer [4]. P53 functions in immunity by induction of apoptosis, removal of apoptotic cells, antiviral defense, induction of type I IFN, enhanced pathogen recognition, cytokine production, and immune checkpoint regulation [4]. Several studies have explored the association of p53 and tumor immune regulation [5–8]. For example, activation of p53 in the tumor microenvironment (TME) might overcome tumor immune suppression and enhance antitumor immunity [8]. P53 could transactivate many tumor immunosuppressive genes such as *PD-L1, VISTA*, and *FOXP3* [5]. P53 functioned in both tumor suppression and anticancer immunosurveillance *via* regulation of *VISTA* [5]. An intriguing question is whether *TP53* mutations are capable of changing the tumor immune microenvironment (TIM), and if yes, what changes may occur in tumor immunity?

Recently, cancer immunotherapy has shown high efficacy in treating a variety of cancers [9]. Particularly, the blockade of immune checkpoints is achieving rapid clinical successes in various advanced malignancies such as melanoma, non-small cell lung cancer, renal cell cancer, Hodgkin’s lymphoma, bladder cancer, head and neck cancer, and the cancers with DNA mismatch-repair (MMR) deficiency [10]. Unfortunately, only a subset of patients can benefit from such therapy [11]. Some molecular biomarkers associated with cancer immunotherapy response have been explored. For example, high tumor mutation burden (TMB) and neoantigen load, MMR deficiency, and expression of PD-1, PD-1 ligands or cytolytic markers in the TIM were associated with positive clinical response to immune checkpoint blockade in cancers [12–17]. However, few studies have correlated the *TP53* mutation status with cancer immunotherapy response, although a recent clinical trial (phase II) data showed that patients with mutated-p53 metastatic breast cancer had better overall survival (OS) prognosis when treated with the immuno-oncology viral agent REOLYSIN^®^ in combination with paclitaxel [18].

Breast cancer (BC) is the most common cancer in women, and is the second leading cause of cancer death in women [19]. About 15-20% of BCs are the triple-negative BC (TNBC), a BC subtype that is clinically negative for expression of the estrogen receptor (ER) and progesterone receptor (PR), and lacks overexpression of the human epidermal growth factor receptor 2 (HER2) [20]. TNBC has a high *TP53* mutation rate (80% in TNBC versus 33% in general BC), and has a poor prognosis due to its aggressive clinical behavior and lack of response to hormonal or HER2 receptor-targeted therapy. Although currently BC does not belong to the cancer types that can be treated by the approved immunotherapeutic drugs, several studies have indicated that TNBC might be propitious to immunotherapy [21, 22].

To explore the association of *TP53* mutations with tumor immunity in BC, we compared the activity of 26 immune cells, function or pathways between *TP53*-mutated and *TP53*-wildtype BCs based on the Cancer Genome Atlas (TCGA) [23] and METABRIC [24] BC genomic data. The 26 immune cells, function or pathways included 15 immune cell types and function, tumor-infiltrating lymphocytes (TILs), immune cell infiltration, regulatory T (Treg) cells, immune checkpoint, cancer testis (CT) antigen, human leukocyte antigen (HLA), cytokine and cytokine receptor (CCR), metastasis-promoting, metastasis-inhibiting, pro-inflammatory, and parainflammation (PI). We found that immune activities in *TP53*-mutated BCs were significantly higher than *TP53*-wildtype BCs. Furthermore, we explored the molecular cues that were associated with the differences in immune activities between *TP53*-mutated and *TP53*-wildtype BCs. Finally, we performed experimental validation of the findings from bioinformatics analysis.

## Methods

### Comparisons of gene expression levels, gene-set enrichment levels, and protein expression levels between two classes of samples

We normalized the TCGA BC gene expression values by base-2 log transformation, and used the original METABRIC gene expression data since they have been normalized. For the TCGA BC protein expression profiles data, we used the downloaded normalized data. We quantified the activity of a cell type, function or pathway in a sample by the single-sample gene-set enrichment analysis (ssGSEA) score (a higher ssGSEA score indicated a higher activity) [25, 26]. We compared expression levels of a single gene or protein between two classes of samples using Student’s *t* test, and compared enrichment levels (ssGSEA scores) of a gene-set or pathway between two classes of samples using Mann-Whitney U test. The false discovery rate (FDR) was used to adjust for multiple tests by the Benjami and Hochberg (BH) method [27]. The threshold of FDR < 0.05 was used to identify the statistical significance. The comparisons involving normal tissue were performed only in TCGA since METABRIC had no normal tissue related data available.

### Comparison of the immune cell infiltration degree between *TP53*-mutated and *TP53*-wildtype BCs

We evaluated the degree **of** immune cell infiltration in the TME in BC using ESTIMATE [28]. For each BC sample, we obtained an immune score to quantify the **d**egree **of** immune cell infiltration in the BC tissue. In addition, we obtained the lymphocyte infiltration percent data for BC from the TCGA BC clinical data. We compared the immune scores or lymphocyte infiltration percents between *TP53*-mutated and *TP53*-wildtype BCs using Mann-Whitney U test.

### Gene-set enrichment analysis

We performed pathway analysis of the set of genes that were differentially expressed between *TP53*-mutated and *TP53*-widtype BCs using the gene-set enrichment analysis (GSEA) software [29]. The threshold of FDR < 0.05 was used to identify the KEGG pathways that had differential activities between *TP53*-mutated and *TP53*-widtype BCs.

### Comparison of proportions of leukocyte cell subsets in the TME between *TP53*-mutated and *TP53*-wildtype BCs

We used CIBERSORT [30] to evaluate the proportions of 22 human leukocyte cell subsets, and compared the proportions of each cell subsets between *TP53*-mutated and *TP53*-wildtype BCs using Mann-Whitney U test. The threshold of FDR < 0.05 was used to identify the leukocyte cell subsets that had significantly different proportions between *TP53*-mutated and *TP53*-wildtype BCs.

### Correlations of pathway or protein activities with immune activities in BC

We obtained the gene-set collections for the p53-mediated pathways from KEGG [31], and quantified the activity of a pathway in a BC sample by the ssGSEA score [25, 26]. To correct for the strong correlation between the p53 pathway and the other p53-mediated pathways, we used the first-order partial correlation to evaluate the correlations between the pathway activity and the immune activity with the R package “ppcor” [32]. The correlation between a pathway activity and an immune activity was defined as significant if FDR<0.05. We quantified the correlation between a protein and an immune gene-set by the Spearman correlation coefficient (“rho”) of expression levels of the protein and ssGSEA scores of the immune gene-set.

### Survival analyses

We compared OS and DFS time between two classes of BC patients classified based on the TP53 mutation, gene expression levels, gene-set enrichment levels, and immune scores, respectively. Kaplan-Meier survival curves were used to show the survival differences between both classes of patients. We classified patients into two different groups based on the TP53 mutation status, or using the median values of gene expression levels, gene-set enrichment levels (ssGSEA scores), or immune scores. The log-rank test was used to calculate the significance of survival-time differences between two classes of patients with a threshold of P < 0.05.

### Comparison of mutation counts between *TP53*-mutated and *TP53*-wildtype BCs

We compared mutation counts between *TP53*-mutated and *TP53*-wildtype BCs using Mann-Whitney U test. This comparison was performed only in TCGA since somatic mutation data in TCGA were generated by whole exome sequencing while in METABRIC they were generated by targeted exome sequencing.

### Cell lines and cell culture

Human cells from breast cancer, MCF-7, and natural killer cells NK-92 were from the American Type Culture Collection (ATCC). MCF-7 was cultured in RPMI-1640 (GIBCO, USA) supplemented with 10% fetal bovine serum (FBS, GIBCO, USA). NK92 cells were incubated in α-MEM (GIBCO, USA) with 2 mM L-glutamine, 0.2 mM inositol, 0.02 mM folic acid, 0.01 mM 2-mercaptoethanol, 10 ng/ml IL-2, 12.5% FBS, and 12.5% horse serum (GIBCO, USA). All the cells were cultured in a humidified incubator at 37◻°C and a 5% CO_2_ atmosphere. Cells were harvested in logarithmic growth phase in all the experiments performed in this study.

### Cell transfection

The MCF-7 cells without antibiotic were maintained in the medium with *TP53*-mutated (c.596 G > T, c.818 A > G, and c.925 T > C) virus stock solution and polybrene (5 ug/mL) for 24h. The transfected MCF-7 cells were cultured in a humidified incubator at 37◻°C and a 5% CO_2_ atmosphere for 48h.

### Co-culture of MCF-7 and NK-92 cells

The transwell chamber (Corning Inc., Corning, NY, USA) was inserted into a 6-well plate to construct a co-culture system. MCF-7 cells were seeded on the 6-well plate at a density of 5×10^4^ cells/well, and NK-92 cells were seeded on the membrane (polyethylene terephthalate, pore size, 0.4μm) of the transwell chamber at a density of 5×10^4^ cells/chamber. NK-92 and MCF-7 cells were co-cultured in a humidified incubator at 37◻°C and a 5% CO_2_ atmosphere for 24h.

### Transwell migration assay

After co-culture of 24h, NK-92 cells were harvested and re-suspended in the upper transwell chambers (8-μm pores, Corning), and MCF-7 cells in the lower 24-well plates. Both NK-92 and MCF-7 cells were incubated at 37 °C for 24h. The membrane was removed and its upper surface was wiped away with a cotton swab to remove the unmigrated NK-92 cells. The membrane was fixed in neutral formalin and air-dried at a room temperature, and was stained with 0.1% crystal violet at 37 °C for 30◻min. The number of NK-92 cells that migrated to the lower surface of the membrane was counted under light microscope. Each assay was performed in triplicate wells.

### EdU proliferation assay

After co-culture of 24h, an EdU (5-ethynyl-2′-deoxyuridine, Invitrogen, CA,USA) proliferation assay [33] was performed to measure the proliferation ability of NK-92 cells. NK-92 cells were plated in 96-well plates at a density of 2◻×◻10^3^ cells/well for 24h. The cells were incubated with 10 μM EdU for 24h at 37 °C before fixation, permeabilization, and EdU staining according to manufacturer’s protocol. The cell nuclei were stained with DAPI (Sigma) at a concentration of 1 μg/ml for 20 min. The proportion of the cells incorporated EdU was determined with fluorescence microscopy. Each assay was performed in triplicate wells.

### CCL4 and CCL5 enzyme-linked immunosorbent assay

After co-culture of 24h, supernatants from NK-92 cells were collected and assayed. CCL4 and CCL5 protein levels were evaluated by ELISA, according to manufacturer’s protocol (Shanghai Enzyme-linked Biotechnology Co., Ltd. China). Study samples and standard dilutions of the chemokines/cytokines were assayed in triplicate. The absorbance was read at 450 nm with the correction set to 570 nm by a microplate reader (BioTek, USA).

### Reverse transcribed quantitative PCR (qPCR) analysis

Z-DEVD-FMK, abemaciclib, and MPP were purchased from Selleck, Haoyuan Chemexpress, and Cayman, respectively. MCF-7 cells were harvested after treated by drugs (Z-DEVD-FMK, 50 μM, 72h; abemaciclib, 250 nM, 7 days; MPP, 0.1 nM, 72h). The total RNA was isolated by Trizol (Invitrogen, USA), and was reverse transcribed into cDNA using the RevertAid First Strand cDNA Synthesis Kit (Thermo Fisher, USA). Primer sequences used for qPCR were presented in the Supplementary Table S16 (Additional file 3). Primers were diluted in nuclease-free water with the Real time PCR Master Mix (SYBR Green) (TOYOBO Co., LTD, JAPAN). Relative copy number was determined by calculating the fold-change difference in the gene of interest relative to β-actin. The qPCR was performed on an ABI 7500 FAST and Applied Biosystems StepOnePlus Real Time PCR machine.

### Western blotting

MCF-7 cells were washed twice with cold PBS and were lysed in SDS buffer (1% SDS, 0.1 M Tris pH 7.4, 10% glycerol) supplemented with protease inhibitors. The protein concentration was determined by Bradford Protein Assay (Bio-rad). After normalization of the total protein content, samples were resolved by standard SDS-PAGE. After Western transfer, NC membranes (Millipore) were incubated with antibodies ER-alpha (21244-1-AP, Proteintech Group, INC.), and cleaved-caspase 3 (KGYC0004-6, KeyGEN Biotech, China). After incubation with the HRP-labeled secondary antibody (KGAA002-1, KeyGEN Biotech, China), proteins were visualized by enhanced chemiluminescence using an G:BOX chemiXR5 digital imaging system.

### Statistical analysis

All experimental data were expressed as mean ± SD, and were analyzed by t test using Prism 5.0 software (GraphPad). P < 0.05 was considered statistically significant.

## Results

### *TP53*-mutated BCs show significantly higher immune activities than *TP53*-wildtype BCs

We quantified the activity of a cell type, function or pathway in a sample by the single-sample gene-set enrichment analysis (ssGSEA) score (a higher ssGSEA score indicated a higher activity) [25, 26]. Strikingly, we found that almost all the immune activities analyzed were significantly higher in *TP53*-mutated BCs than in *TP53*-wildtype BCs consistently in both TCGA and METABRIC datasets (Mann-Whitney U test, P<0.05; Figure 1A; Additional File 4, Figure S1A). Moreover, *TP53*-mutated BCs had significantly higher immune scores than *TP53*-wildtype BCs in both datasets (Mann-Whitney U test, P=1.35*10^−7^, 1.75*10^−34^ for TCGA and METABRIC, respectively) (Figure 1B). Based on the TCGA BC pathological slides data, we found that *TP53*-mutated BCs had markedly higher percent of lymphocyte infiltration compared to *TP53*-wildtype BCs (Mann-Whitney U test, P=0.01) (Figure 1C). Altogether, these data indicated that *TP53* mutations were associated with elevated immune activities in BC.

**Figure 1.**
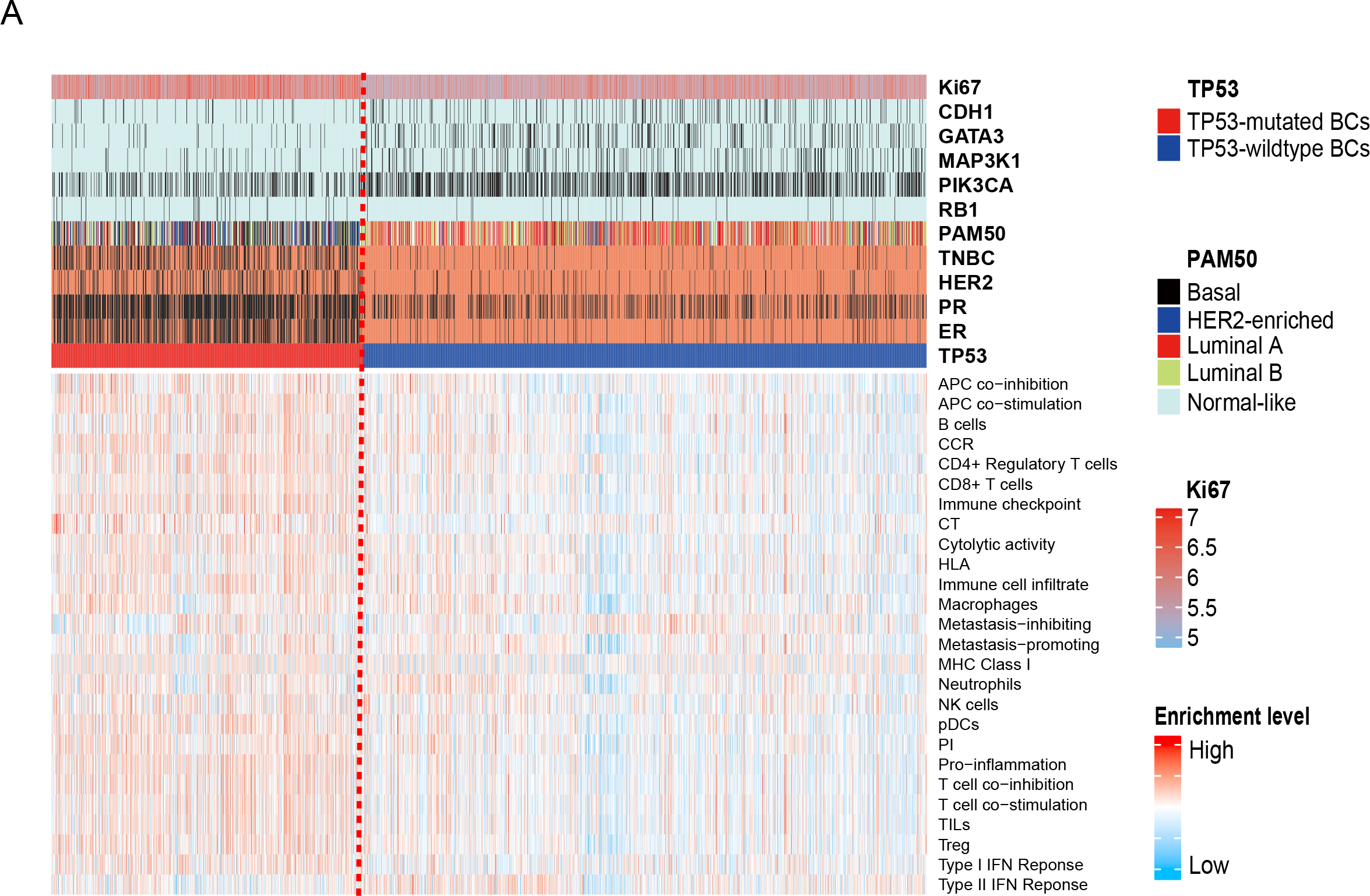
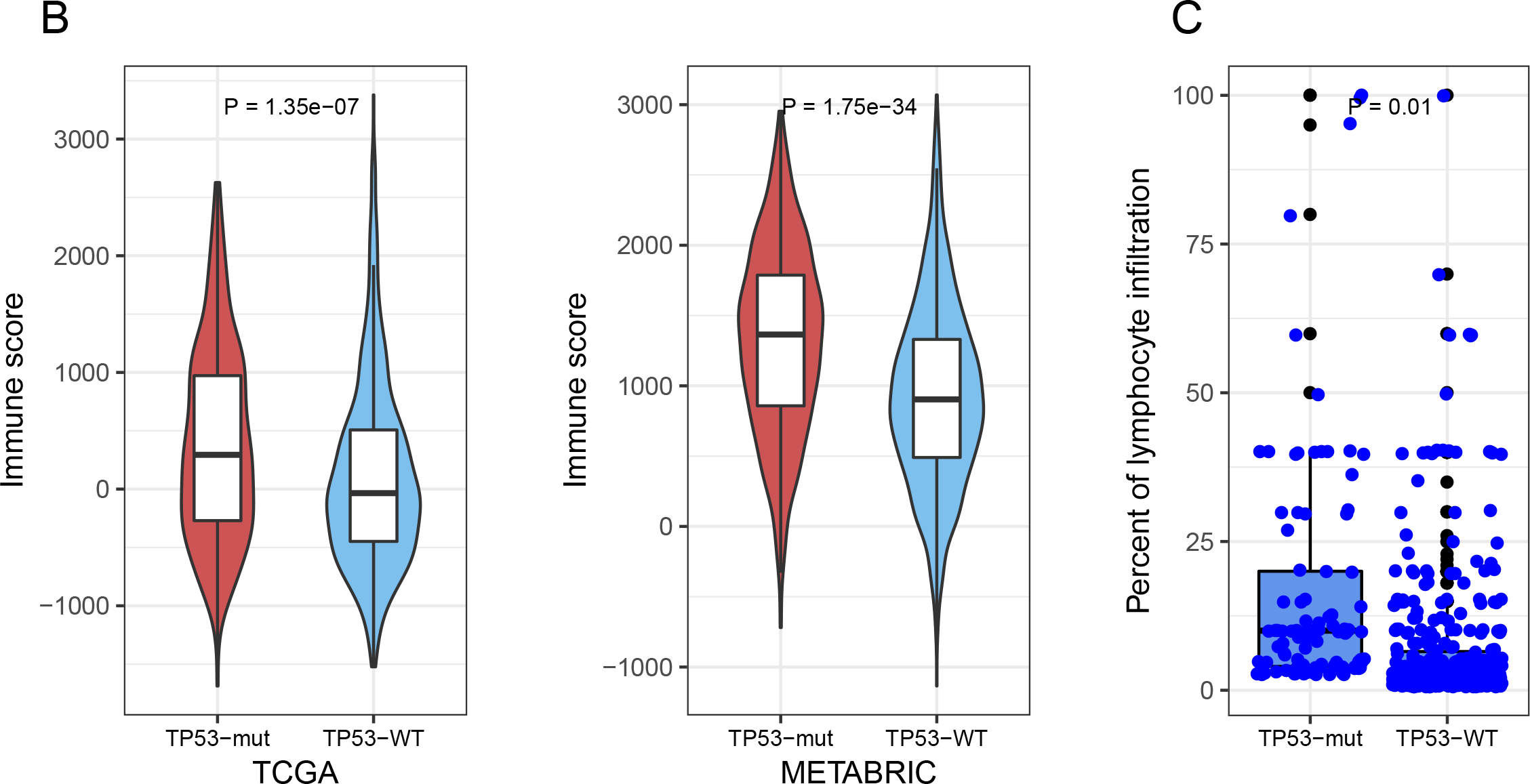
*TP53*-mutated breast cancers (BCs) have increased immune activities compared to *TP53*-wildtype BCs. A. Heatmap showing the ssGSEA scores of 26 immune gene-sets in *TP53*-mutated and *TP53*-wildtype BCs (METABRIC). ssGSEA: single-sample gene-set enrichment analysis. TNBC: triple-negative breast cancer. *RB1* are more frequently mutated in *TP53*-mutated BCs while *CDH1, GATA3, MAP3K1*, and *PIK3CA* are more frequently mutated in *TP53*-wildtype BCs (Fisher’s exact test, P<0.05). The black vertical lines in the horizontal bars beside gene symbols indicate that the genes are mutated in corresponding samples. The black vertical lines in the horizontal bar beside “TNBC” indicate that the sample is a TNBC. The black vertical lines in the horizontal bars beside “ER”, “PR”, and “HER2” indicate that the sample is ER-, PR-, or HER2-. B. *TP53*-mutated BCs have significant higher degree of immune infiltration than *TP53*-wildtype cancers evaluated by ESTIMATE [28]. C. The TCGA BC pathological slides data show that *TP53*-mutated BCs had markedly higher percent of lymphocyte infiltration than *TP53*-wildtype BCs. *TP53*-mut: *TP53*-mutated BCs. *TP53*-WT: *TP53*-wildtype BCs. It applies to all the other figures.

### *TP53* mutations are associated with elevated expression of immune cell types and functional marker genes in BC

We analyzed 15 immune cell types and functional gene-sets that were associated with B cells, CD4+ regulatory T cells, CD8+ T cells, macrophages, neutrophils, natural killer (NK) cells, plasmacytoid dendritic cells (pDCs), histocompatibility complex (MHC) class I, antigen-presenting cell (APC) costimulation, T cell co-stimulation, APC co-inhibition, T cell co-inhibition, Type I IFN response, TypeII IFN response, and cytolytic activity, respectively [34]. We found that a substantial number of genes in the 15 gene-sets had significantly higher expression levels in *TP53*-mutated BCs than in *TP53*-wildtype BCs in both TCGA and METABRIC datasets (Figure 2A; Additional file 4, Figure S1B). For example, 12 of the 13 T cell co-stimulation marker genes, the CD8+ T cell marker gene (*CD8A*), – the MHC Class I marker genes (*B2M, HLA-A*, and *TAP1*), and the cytolytic activity genes (*GZMA* and *PRF1*) were more highly expressed in *TP53*-mutated BCs than in *TP53*-wildtype BCs (Additional file 1, Table S1). Moreover, 14 of the 15 gene-sets showed significantly higher enrichment levels in *TP53*-mutated BCs than in *TP53*-wildtype BCs in at least one dataset, and 12 in both datasets (Mann-Whitney U test, FDR<0.05) (Additional file 1, Table S1; Additional file 4, Figure S1C). These results suggest that *TP53*-mutated BCs likely have elevated immunogenic activity compared to *TP53*-wildtype BCs.

**Figure 2.**
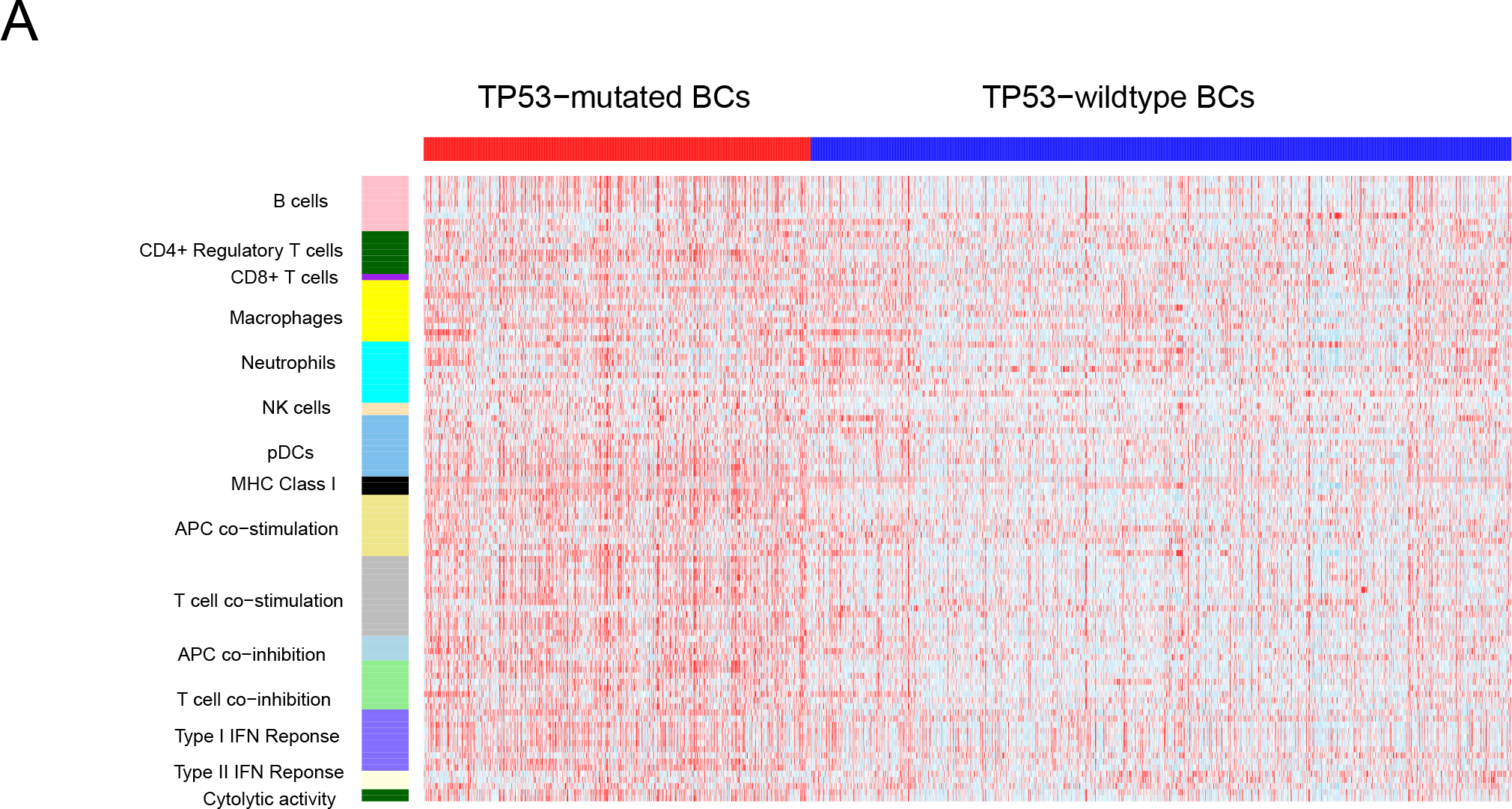
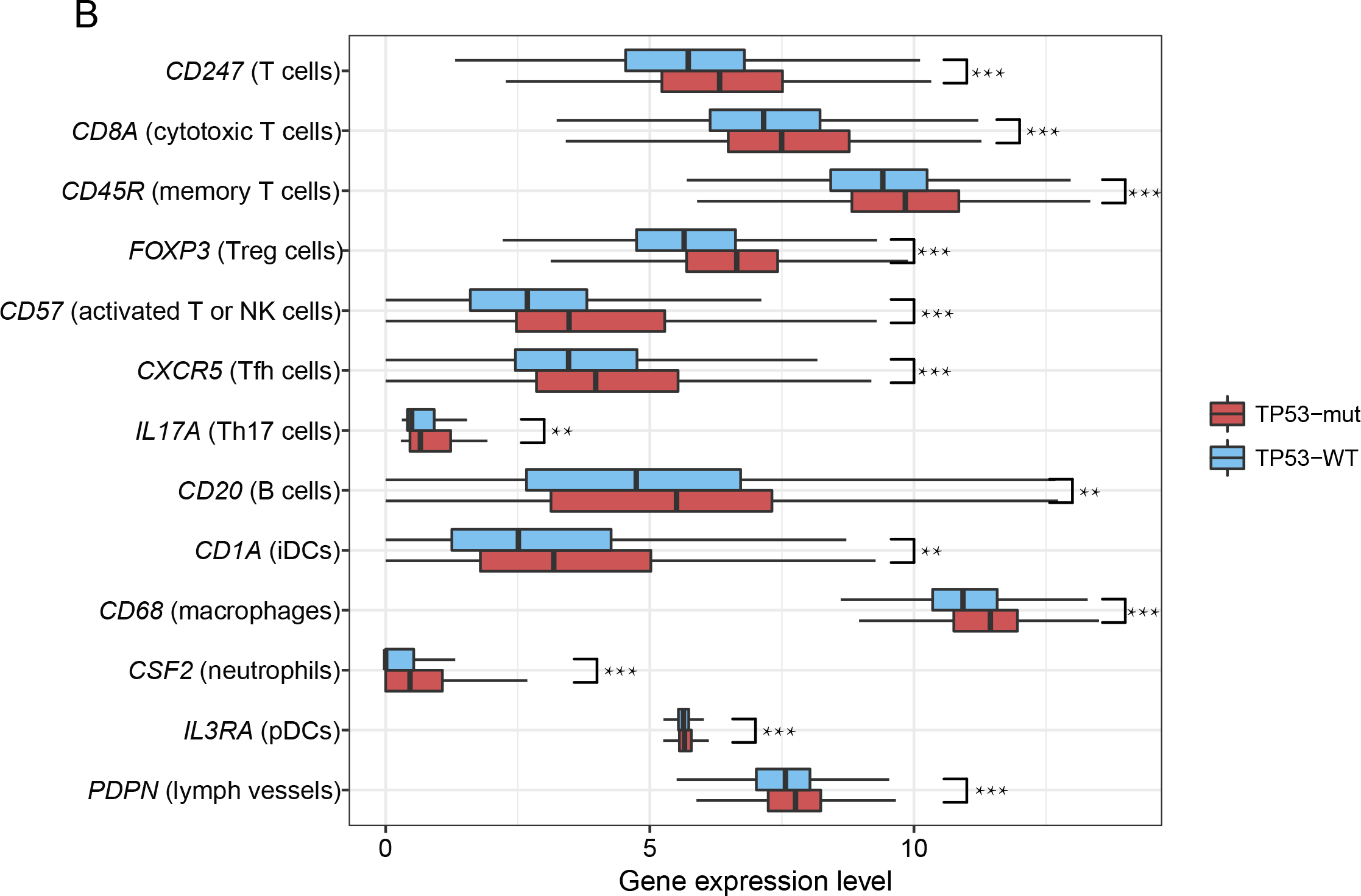
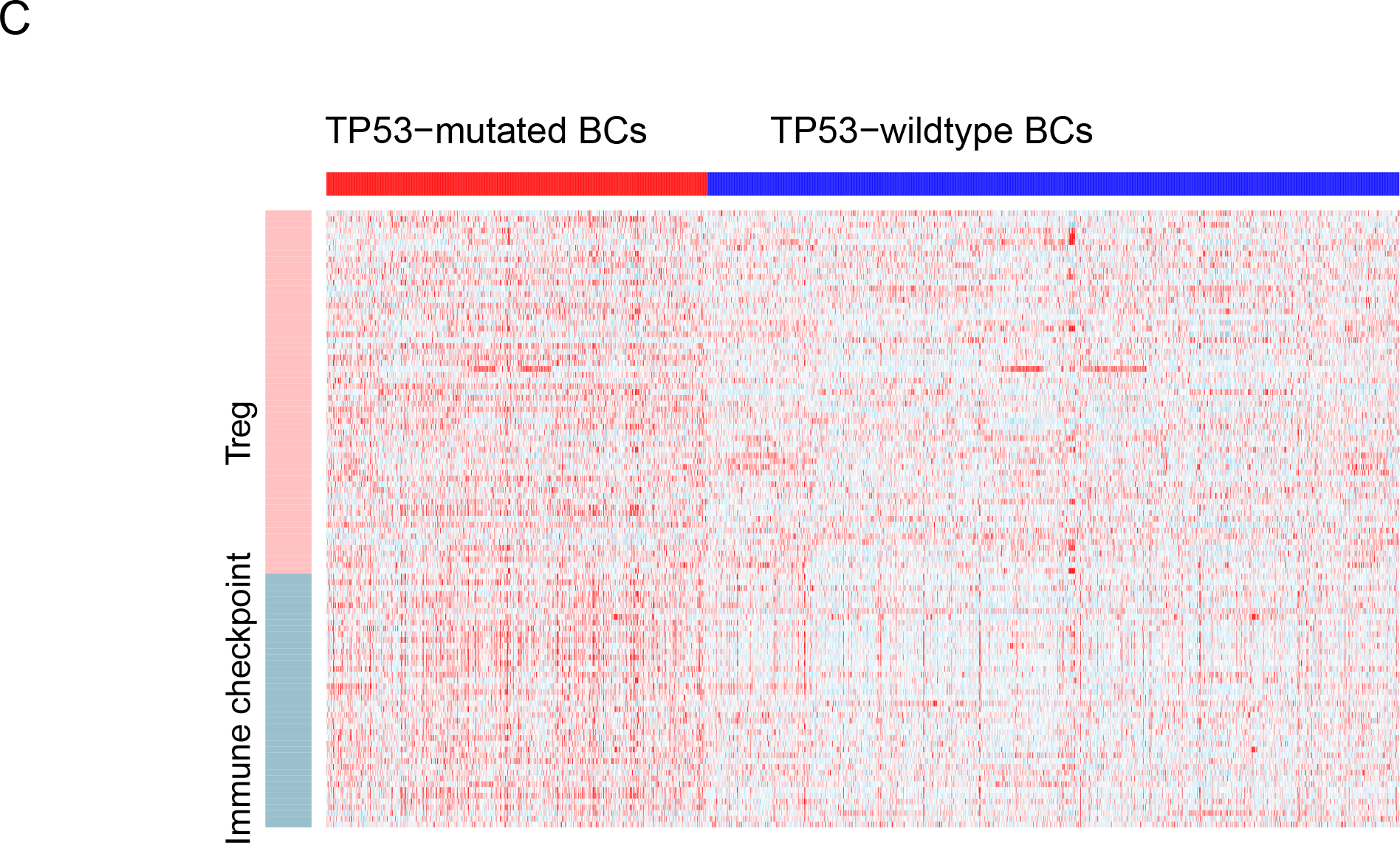
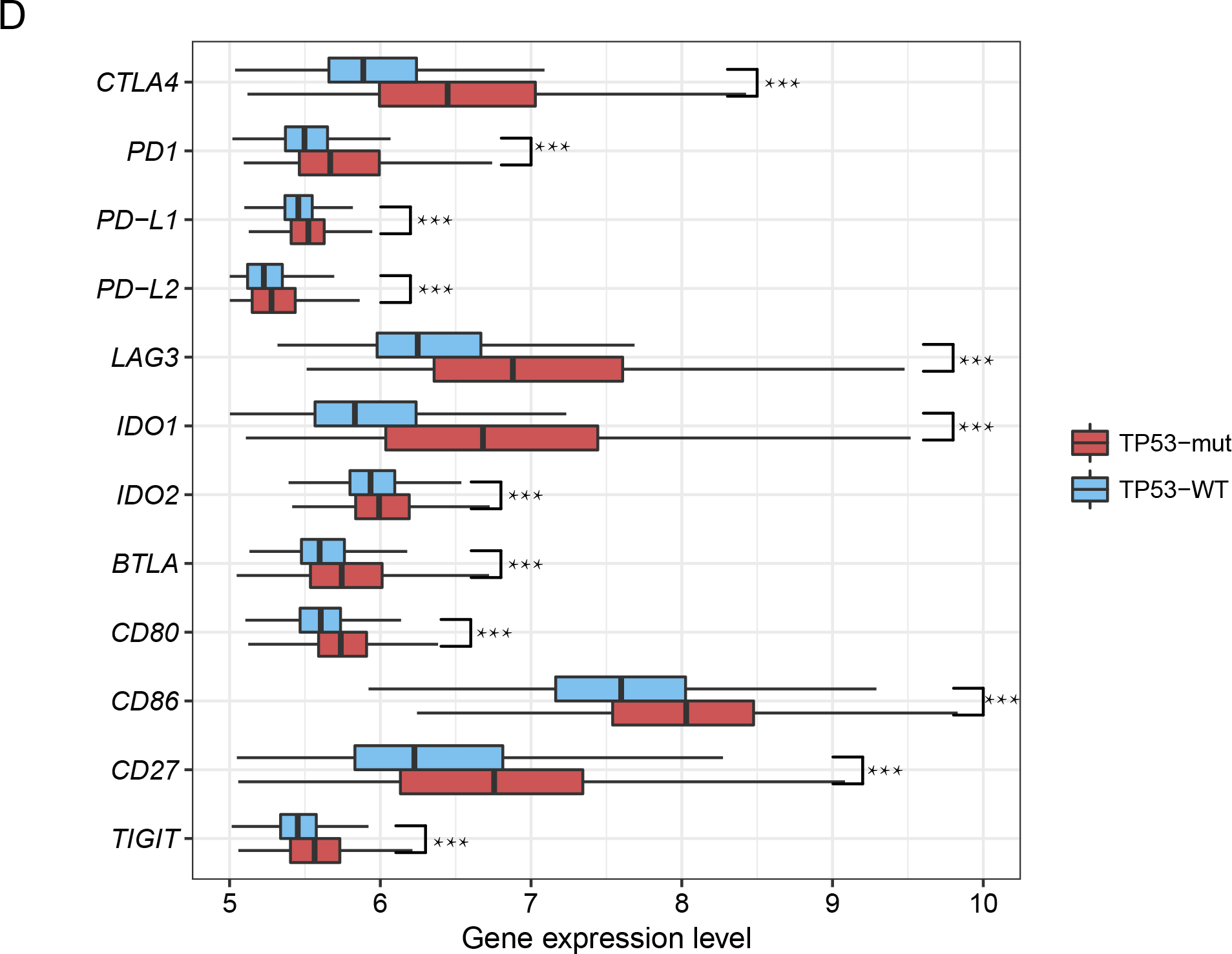
*TP53*-mutated breast cancers (BCs) likely have elevated expression of immune cell types and functional marker, Treg, and immune checkpoint genes compared to *TP53*-wildtype BCs. A. Heatmap showing *TP53*-mutated BCs likely more highly express immune cell types and functional marker genes than *TP53*-wildtype BCs (METABRIC). NK: natural killer. pDCs: plasmacytoid dendritic cells. MHC: histocompatibility complex. APC: antigen-presenting cell. B. Immune cell subpopulations marker genes likely have higher expression levels in *TP53*-mutated BCs than in *TP53*-wildtype BCs. C. Heatmap showing *TP53*-mutated BCs likely more highly express Treg and immune checkpoint genes than *TP53*-wildtype BCs (METABRIC). D. A number of important immune checkpoint genes are upregulated in *TP53*-mutated BCs versus *TP53*-wildtype BCs. Treg: regulatory T cell. Red color indicates higher gene expression levels, and blue color indicates lower gene expression levels in the heatmaps. *: P < 0.05; **: P < 0.01; ***: P < 0.001, and it applies to all the other figures.

### *TP53* mutations are associated with higher degree of immune cell infiltration in BC

We compared expression levels of 122 TILs gene signatures [35] between *TP53*-mutated and *TP53*-wildtype BCs. Strikingly, 112 (92%) TILs genes were more highly expressed in *TP53*-mutated BCs in at least one dataset (88 in both datasets) (Additional file 1, Table S2; Additional file 4, Figure S1D). The ssGSEA scores for the TILs gene-set were significantly higher in *TP53*-mutated BCs than in *TP53*-wildtype BCs (Mann-Whitney U test, P=2.84*10^−8^, 4.22*10^−33^ for TCGA and METABRIC, respectively) (Additional file 4, Figure S1E). Compared to normal tissue, the ssGSEA scores for the TILs gene-set were significantly higher in *TP53*-mutated BCs (Mann-Whitney U test, P=0.001), while had no significant differences in *TP53*-wildtype BCs (Mann-Whitney U test, P=0.18). These data indicate that *TP53*-mutated BCs have higher degree of TILs infiltration than *TP53*-wildtype BCs.

In addition, we compared the immune infiltrate densities of 15 different immune cell subpopulations between *TP53*-mutated and *TP53*-wildtype BCs. These immune cell subpopulations included T cells (quantified with marker *CD247* gene expression levels), cytotoxic T cells (*CD8A*), memory T cells (*CD45R*), Tregs (*FOXP3*), activated T or NK cells (*CD57*), Tfh cells (*CXCR5*), Th17 cells (*IL-17*), B cells (*CD20*), immature dendritic cells (iDCs) (*CD1A*), pDCs (*IL3RA*), macrophages (*CD68*), mast cells (*TPSAB1*), neutrophils (*CSF2*), blood vessels (*ENG*), and lymph vessels (*PDPN*) [36]. Strikingly, 13 of the 15 immune cell subpopulation marker genes had significantly higher expression levels in *TP53*-mutated BCs in a single or both datasets (Figure 2B; Additional file 1, Table S2). Interestingly, compared to normal tissue, *CD57, ENG, IL3RA*, and *PDPN* were more lowly expressed in both *TP53*-mutated and *TP53*-wildtype BCs, while *CD1A, FOXP3*, and *CXCR5* were more highly expressed in both BC subtypes. The decreased subpopulations of activated T cells, NK cells, and pDCs, and increased subpopulations of Treg cells, iDCs, and Tfh cells in BC may suggest an immune evasion mechanism in BC [37].

### *TP53* mutations are associated with higher immunosuppressive activity in BC

Treg cells are crucial for the maintenance of immunosuppressive activity in cancer [38], and immune checkpoints play an important role in tumor immunosuppression [39, 40]. We compared expression levels of 70 Treg gene signatures and 47 immune checkpoint genes [41] between *TP53*-mutated and *TP53*-wildtype BCs, respectively. We found that 43 (61%) Treg and 40 (85%) immune checkpoint genes were more highly expressed in *TP53*-mutated BCs in a single or both datasets, respectively (Figure 2C; Additional file 1, Table S3, S4; Additional file 4, Figure S1F). Moreover, *TP53*-mutated BCs had remarkably higher enrichment levels of the Treg and immune checkpoint gene-sets than *TP53*-wildtype BCs in both datasets (Mann-Whitney U test, P<10^−10^) (Additional file 4, Figure S1G). Interestingly, a number of immune checkpoint genes upregulated in *TP53*-mutated BCs have been established or considered promising targets for cancer immunotherapy including *CTLA4, PD1, PD-L1, PD-L2, LAG3, IDO1/2, BTLA, CD80, CD86, CD27*, and *TIGIT* (Figure 2D; Additional file 1, Table S4; Additional file 4, Figure S1H). All these genes except *PD-L1* were also upregulated in *TP53*-mutated BCs compared to normal tissue.

Altogether, these results showed that *TP53*-mutated BCs likely had higher activities of Treg cell infiltration and the immune checkpoint pathway than *TP53*-wildtype BCs. It suggests that p53 may play a role in inhibiting tumor immunosuppression in BC.

### *TP53* mutations are associated with higher cytokine activity in BC

Cytokines are a group of small proteins that are important components within the TME, and play important roles in tumor immunity [42]. We compared expression levels of 261 cytokine and cytokine receptor (CCR) genes [43] between *TP53*-mutated and *TP53*-wildtype BCs. We found that the number of CCR genes (158 in a single or both datasets) more highly expressed in *TP53*-mutated BCs far exceeded the number of CCR genes (47 in a single or both datasets) more highly expressed in *TP53*-wildtype BCs (Fisher’s exact test, P<2.2*10^−16^, OR=11.48) (Additional file 1, Table S5; Additional file 5, Figure S2A). Moreover, we found that 230 of the 261 CCR genes showed more frequent somatic copy number alterations (SCNAs) in *TP53*-mutated BCs compared to *TP53*-wildtype BCs (Fisher’s exact test, FDR<0.05; Figure 3A). The ssGSEA scores for the CCR gene-set were significantly higher in *TP53*-mutated BCs than in *TP53*-wildtype BCs in both datasets (Mann-Whitney U test, P=1.9*10^−10^, 5.32*10^−39^ for TCGA and METABRIC, respectively) (Additional file 5, Figure S2B). However, compared to normal tissue, the ssGSEA scores for the CCR gene-set were significantly lower in both *TP53*-mutated BCs and *TP53*-wildtype BCs (Mann-Whitney U test, P=0.04, 5.43*10^−13^ for TCGA and METABRIC, respectively). It indicates that the cytokine activity is downregulated in BC while *TP53*-mutations may promote the cytokine activity in BC.

**Figure 3.**
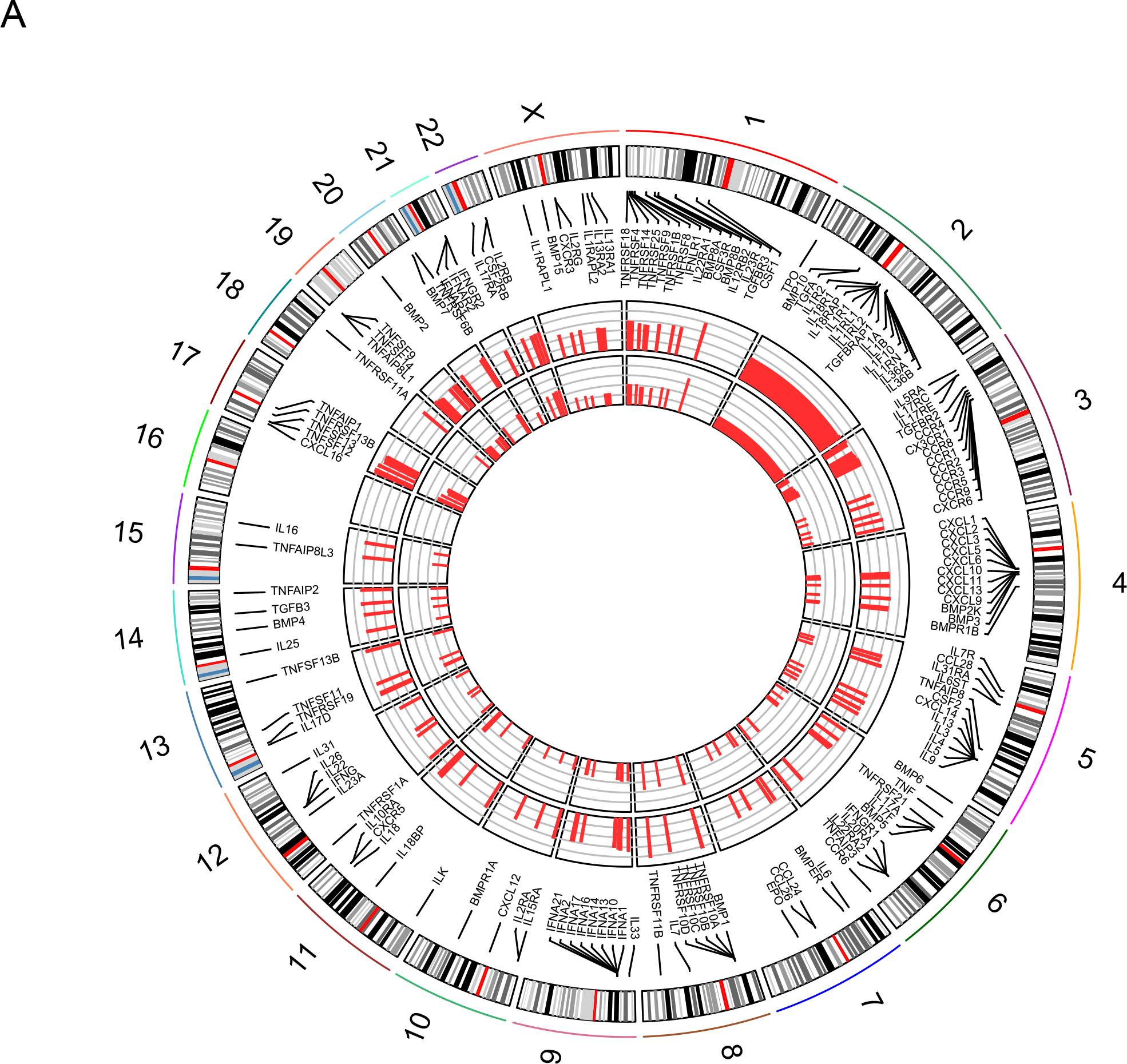
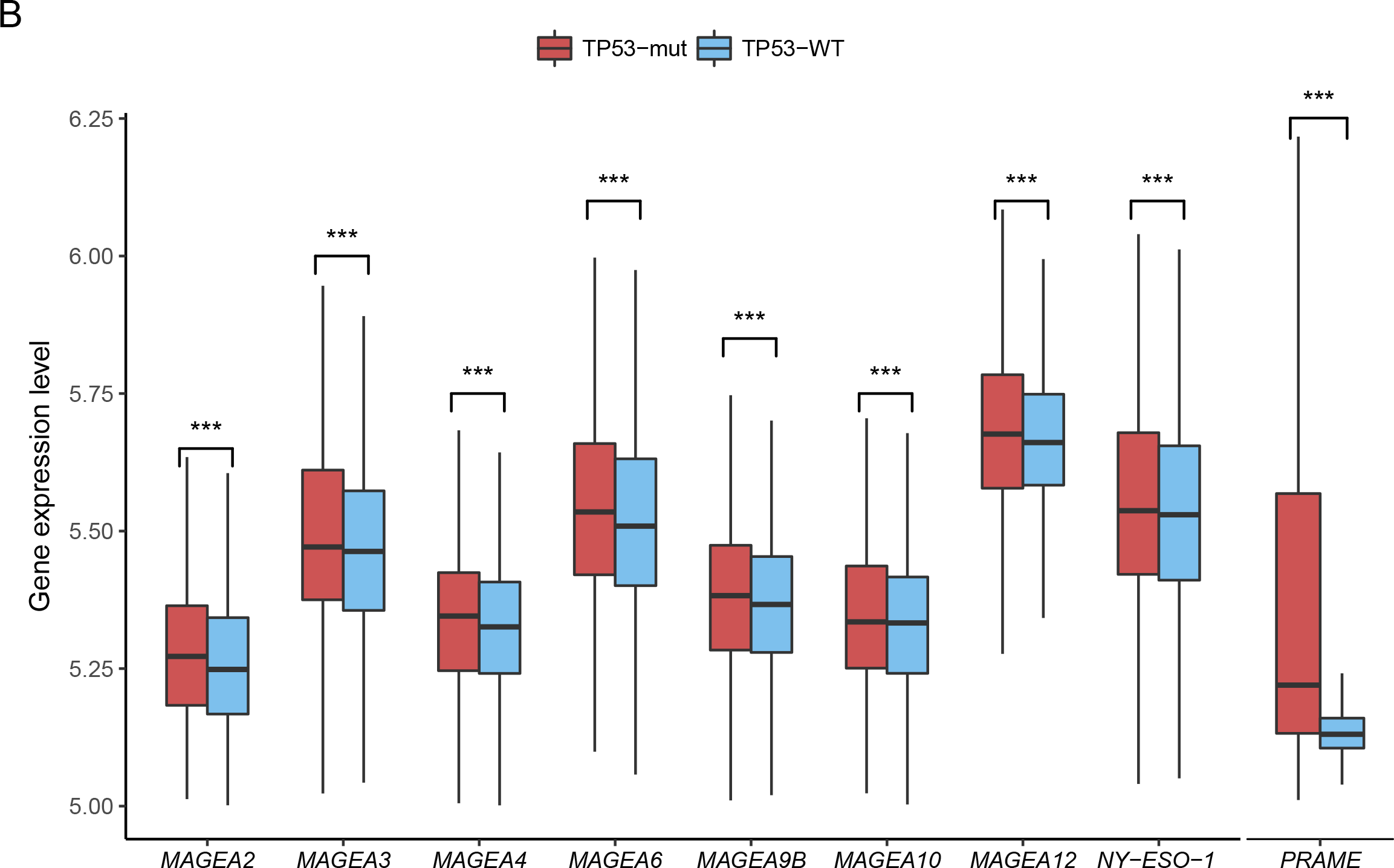
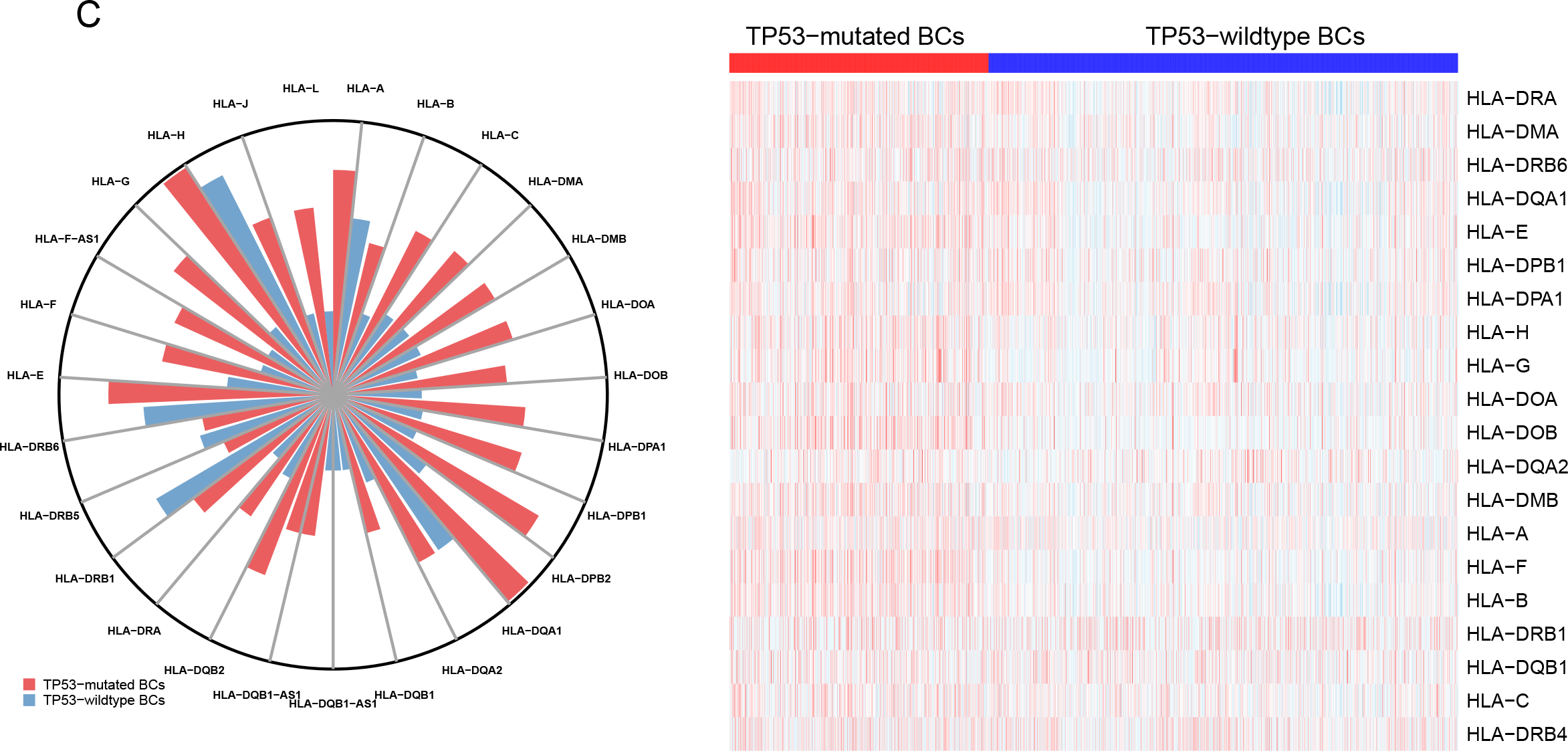
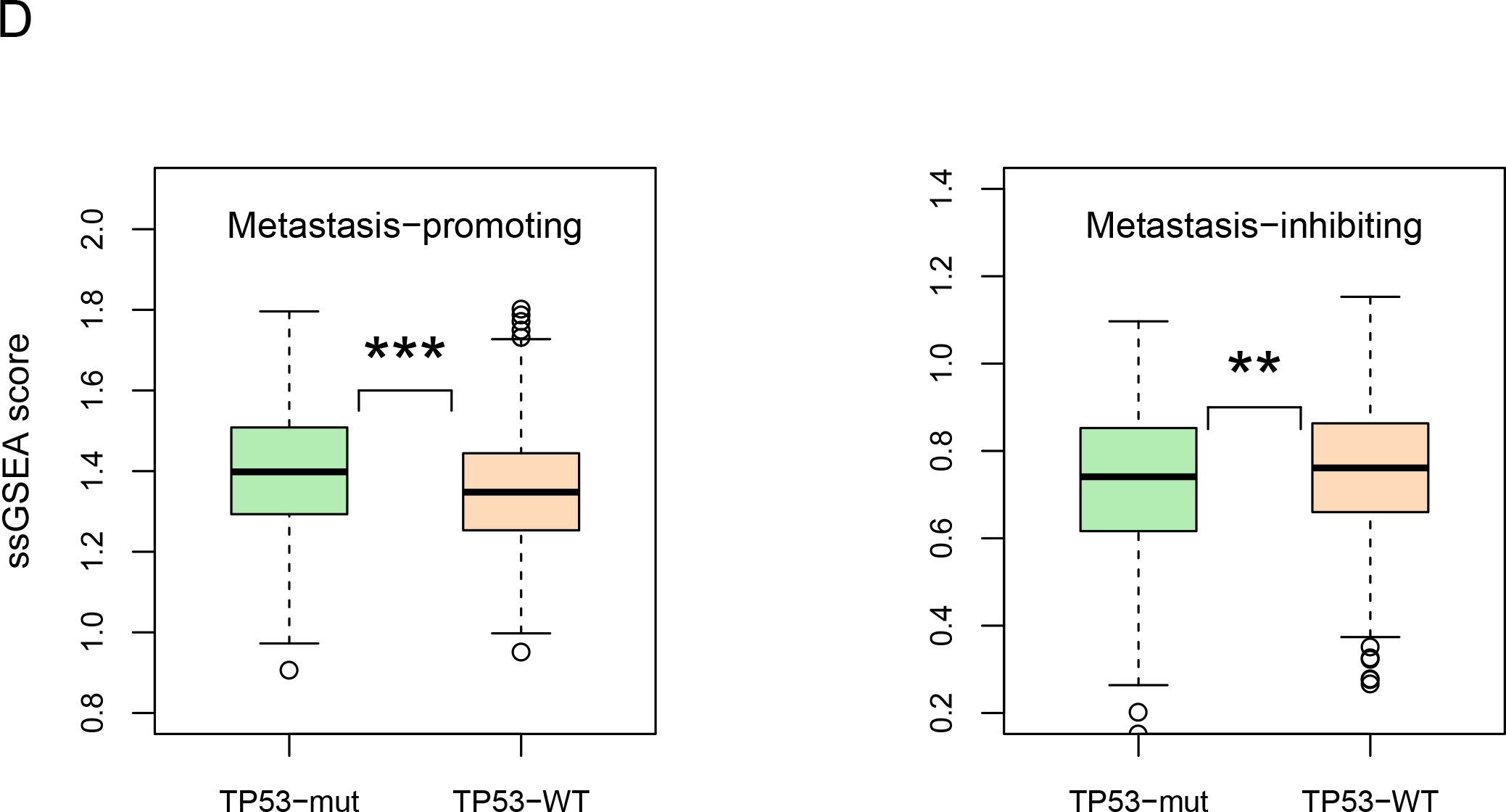
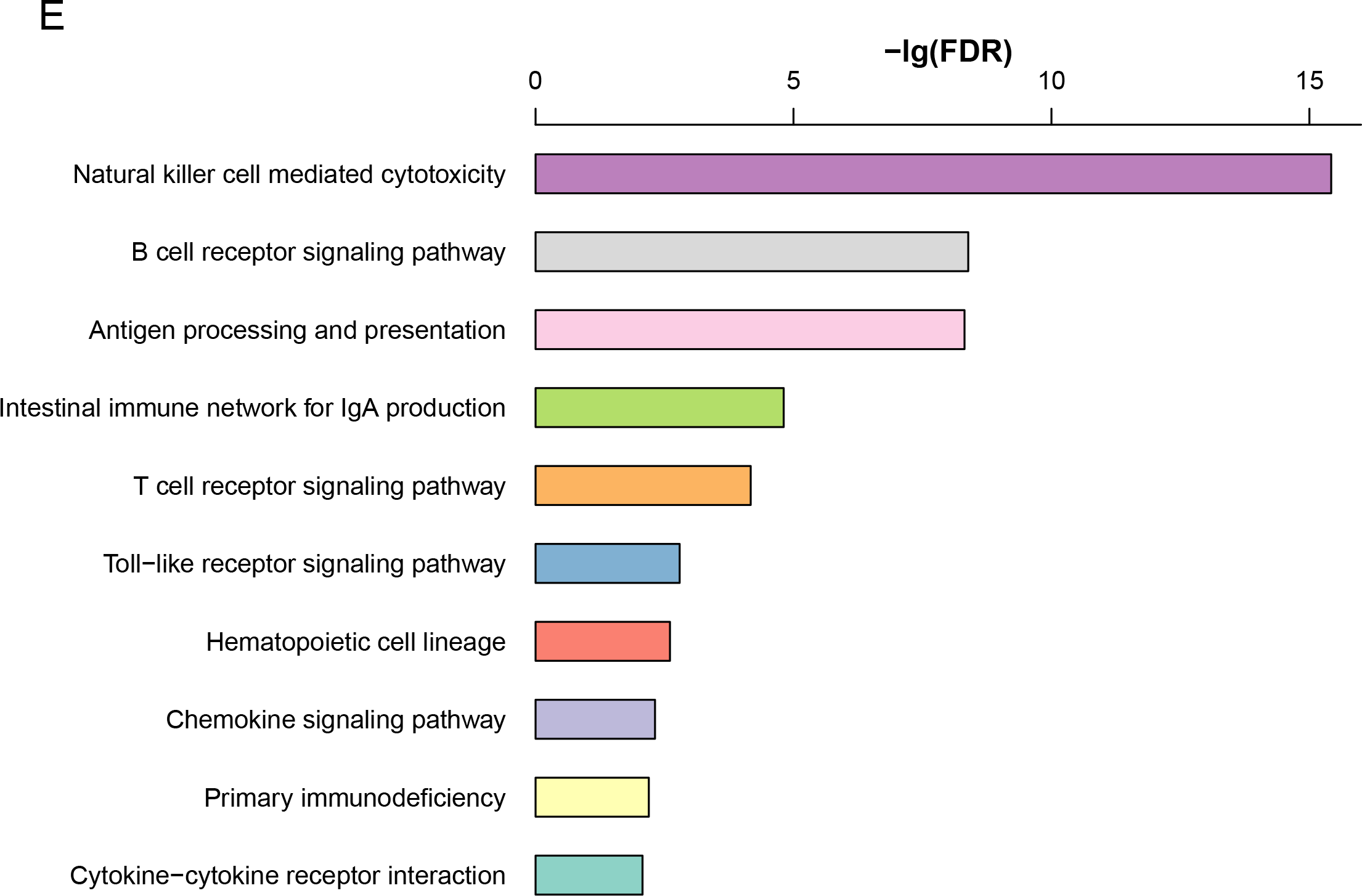
*TP53*-mutated breast cancers (BCs) have higher immune activities of cytokine, cancer-testis antigen, HLA genes, metastasis-promoting, and a number of immune-related KEGG pathways than *TP53*-wildtype BCs. A. Cytokine and cytokine receptor (CCR) genes showing more frequent somatic copy number alterations (SCNAs) in *TP53*-mutated BCs than in *TP53*-wildtype BCs (Fisher’s exact test, FDR<0.05). The outmost circle indicates 23 human chromosomes. The bars in both inner circles (outside and inside) indicate the frequency of SCNAs of CCR genes in *TP53*-mutated and *TP53*-wildtype BCs, respectively. A longer bar indicates a higher frequency of SCNAs. B. Upregulated cancer-testis (CT) antigen genes in *TP53*-mutated BCs versus *TP53*-wildtype BCs encoding the CT antigens that are potential targets for developing cancer vaccines. C. HLA genes are more frequently amplified and more highly expressed in *TP53*-mutated BCs than in *TP53*-wildtype BCs. The length of the bars in the rose diagram is proportional to the frequency of HLA gene amplification in *TP53*-mutated or *TP53*-wildtype BCs. Red color indicates higher gene expression levels, and blue color indicates lower gene expression levels in the right heatmap. D. Enrichment levels of the metastasis-promoting gene-set are significantly higher, and enrichment levels of the metastasis-inhibiting gene-set are significantly lower in *TP53*-mutated BCs than in *TP53*-wildtype BCs (Mann-Whitney U test, P<0.05). E. Immune-related KEGG pathways upregulated in *TP53*-mutated BCs relative to *TP53*-wildtype BCs (FDR q-value<0.05).

### *TP53* mutations are associated with elevated expression of many cancer-testis antigens in BC

Cancer-testis (CT) antigens are a group of immunogenic proteins that are normally expressed only in male germ cells, but are also aberrantly activated in a variety of cancers [44]. We compared expression levels of 233 CT antigen genes [45] between *TP53*-mutated and *TP53*-wildtype BCs. We found 128 CT antigen genes more highly expressed in *TP53*-mutated BCs versus 41 more highly expressed in *TP53*-wildtype BCs in a single or both datasets (Fisher’s exact test, P<2.2*10^−16^, odds ratio (OR)=5.7) (Additional file 1, Table S6). Many of the upregulated CT antigen genes in *TP53*-mutated BCs encode the CT antigens that are potential targets for developing cancer vaccines, including *MAGEA, NY-ESO-1*, and *PRAME* (Figure 3B; Additional file 5, Figure S2C). The enrichment levels of the CT antigen gene-set were significantly higher in *TP53*-mutated BCs than in *TP53*-wildtype BCs in both datasets (Mann-Whitney U test, P=9.52*10^−35^, 9.46*10^−24^ for TCGA and METABRIC, respectively) (Additional file 5, Figure S2D). These results indicated that p53 might repress expression of many CT antigen genes, a finding in line with a prior study showing that p53 regulated CT antigen genes [46]. As expected, the enrichment levels of the CT antigen gene-set were significantly higher in both *TP53*-mutated BCs and *TP53*-wildtype BCs compared to normal tissue (Mann-Whitney U test, P=4.23*10^−53^, 1.8*10^−51^ for TCGA and METABRIC, respectively).

### *TP53* mutations are associated with significant alterations of HLA genotypes and phenotypes in BC

HLA genes encode MHC proteins, which play important roles in the regulation of the immune system [47]. We found that HLA genes were more frequently amplified in *TP53*-mutated BCs compared to *TP53*-wildtype BCs (Figure 3C). Moreover, *TP53*-mutated BCs had lower somatic mutation rates of HLA genes than *TP53*-wildtype BCs in TCGA (Fisher’s exact test, P=0.02, OR=0.6), while METABRIC had no somatic mutation data available for HLA genes. Gene mutations may yield neoepitopes that can be recognized by immune cells [48]. We found that *TP53*-mutated BCs had markedly higher total mutation counts than *TP53*-wildtype BCs in TCGA (Mann-Whitney U test, P=4.18*10^−25^). However, the numbers of gene mutations yielding predicted HLA-binding peptides [34] showed no significant differences between *TP53*-mutated and *TP53*-wildtype BCs (Mann-Whitney U test, P=0.4). It suggests that TMB is not the essential factor explaining the differential immune activities between *TP53*-mutated and *TP53*-wildtype BCs. Furthermore, we found that most HLA genes showed significantly higher expression levels in *TP53*-mutated BCs than in *TP53*-wildtype BCs (Figure 3C; Additional file 1, Table S7; Additional file 6, Figure S3A). These results indicated that *TP53* mutations might promote HLA activity in BC. This finding appears not to be consistent with a previous study showing that p53 increased expression of MHC proteins in cancer [49]. The inconsistency supports the notion that the p53 function is context-dependent, and largely depends on the cell type [50, 51].

### *TP53* mutations are associated with elevated inflammatory activity in BC

Inflammatory responses play important roles in tumor development and often has protumorigenic effects [52]. We compared expression levels of 16 pro-inflammatory genes [53] between *TP53*-mutated and *TP53*-wildtype BCs. Strikingly, all 16 genes had significantly higher expression levels in *TP53*-mutated BCs than in *TP53*-wildtype BCs in at least one dataset (13 in both datasets) (Additional file 6, Figures S3B, S3C; Additional file 1, Table S8). Notably, *STAT1* (signal transducer and activator of transcription 1) was more highly expressed in *TP53*-mutated BCs than in both *TP53*-wildtype BCs and normal tissue. This gene has been shown to interact with p53 [54], and play an important role in maintaining an immunosuppressive TME in BC [55]. Another gene *GZMB* (granzyme B), together with aforementioned cytolytic activity marker gene *GZMA*, were upregulated in *TP53*-mutated BCs compared to both *TP53*-wildtype BCs and normal tissue. The products of both genes are mainly secreted by NK cells and cytotoxic T lymphocytes, and are associated with immune cytolytic activity [34]. Thus, elevated expression of both genes suggested that *TP53* mutations might promote inflammatory and immune cytolytic activities in BC. The ssGSEA scores for the pro-inflammatory gene-set were significantly higher in *TP53*-mutated BCs than in *TP53*-wildtype BCs in both datasets (Mann-Whitney U test, P=2.07*10^−17^, 1.81*10^−61^ for TCGA and METABRIC, respectively).

Parainflammation (PI) is a low-grade inflammatory reaction that plays a role in carcinogenesis [56]. The enrichment levels of the PI gene-set [56] were significantly higher in *TP53*-mutated BCs than in *TP53*-wildtype BCs (Mann-Whitney U test, P=1.03*10^−9^, 5.03*10^−37^ for TCGA and METABRIC, respectively), and were also significantly higher in both *TP53*-mutated BCs and *TP53*-wildtype BCs than in normal tissue (Mann-Whitney U test, P=6.89*10^−12^, 0.003 for *TP53*-mutated and *TP53*-wildtype BCs, respectively) (Additional file 1, Table S8). These data indicated that *TP53* mutations positively correlated with the PI activity in BC, a finding consistent with a previous study showing that PI was associated with the p53 status in cancer [56].

### *TP53* mutations promote BC metastasis *via* immune regulation

In a recent study, Weyden *et al.* identified 23 genes that were involved in immune regulation of tumor metastasis [57]. Among the 23 genes, 15 (*GRSF1, BC017643, CYBB, FAM175B, BACH2, NCF2, ARHGEF1, FBXO7, TBC1D22A, ENTPD1, LRIG1, HSP90AA1, CYBA, NBEAL2*, and *SPNS2*) promoted tumor metastasis, and 8 (*IRF1, RNF10, PIK3CG, DPH6, SLC9A3R2, IGHM, IRF7*, and *ABHD17A*) inhibited tumor metastasis. We found that 9 of the 15 metastasis-promoting genes had higher expression levels in *TP53*-mutated BCs than in *TP53*-wildtype BCs in a single or both datasets (Additional file 1, Table S9). Notably, *SPNS2* (sphingolipid transporter 2) which promoted tumor metastasis *via* regulation of lymphocyte trafficking [57], had higher expression levels in *TP53*-mutated BCs than in *TP53*-wildtype BCs in TCGA (Student’s *t* test, FDR=1.34*10^−6^) while its expression data were lacking in METABRIC. The enrichment levels of the metastasis-promoting gene-set were significantly higher in *TP53*-mutated BCs than in *TP53*-wildtype BCs in both datasets (Mann-Whitney U test, P=3.62*10^−6^, 0.003 for TCGA and METABRIC, respectively) (Figure 3D). In contrast, of the 7 metastasis-inhibiting genes (*IGHM* was excluded since it had no gene expression data available in either of both datasets), 5 had lower expression levels in *TP53*-mutated BCs than in *TP53*-wildtype BCs in a single or both datasets (Additional file 1, Table S9). The metastasis-inhibiting gene-set showed significantly lower enrichment levels in *TP53*-mutated BCs than in *TP53*-wildtype BCs in TCGA (Mann-Whitney U test, P=0.01) (Figure 3D). Altogether, these results suggest that *TP53*-mutated BCs likely have increased expression of metastasis-promoting genes and depressed expression of metastasis-inhibiting genes compared to *TP53*-wildtype BCs, indicating that *TP53*-mutated BCs are metastasis-prone and this characteristic may be associated with the dysfunction of p53 immune response regulation in the TME.

### A number of immune-related pathways are upregulated in *TP53*-mutated BCs relative to *TP53*-wildtype BCs

We performed gene-set enrichment analysis of the set of genes that were upregulated in *TP53*-mutated BCs, and the set of genes that were downregulated in *TP53*-mutated BCs compared to *TP53*-wildtype BCs concurrently in both datasets, respectively (Student’s *t* test, FDR<0.05). We found that a number of immune-related KEGG [31] pathways were significantly associated with the set of genes upregulated in *TP53*-mutated BCs (FDR q-value<0.05). These pathways included natural killer cell mediated cytotoxicity, B cell receptor signaling, antigen processing and presentation, intestinal immune network for IgA production, T cell receptor signaling, toll-like receptor signaling, hematopoietic cell lineage, chemokine signaling, primary immunodeficiency, and cytokine-cytokine receptor interaction (Figure 3E). In contrast, only the cytokine-cytokine receptor interaction pathway was significantly associated with the set of genes downregulated in *TP53*-mutated BCs. These results again demonstrate that *TP53* mutations are associated with elevated immune activities in BC.

### *TP53* mutations are associated with differential immune cell subset infiltration in BC

We compared the proportions of 22 human leukocyte cell subsets that were evaluated by CIBERSORT [30] between *TP53*-mutated and *TP53*-wildtype BCs. We found that activated dendritic cells, M0 macrophages, M1 macrophages, activated T cells CD4 memory, and T cells follicular helper cell subsets had significantly higher proportions in *TP53*-mutated BCs in both datasets (Mann-Whitney U test, FDR<0.05; Figure 4). In contrast, resting dendritic cells, M2 macrophages, resting mast cells, monocytes, and resting T cells CD4 memory cell subsets had significantly higher proportions in *TP53*-wildtype BCs in both datasets (Mann-Whitney U test, FDR<0.05; Figure 4). This further demonstrates that *TP53* mutations are associated with stronger activity of immune cells in BC. Interestingly, M1 macrophages that incite inflammation had higher proportions in *TP53*-mutated BCs than in *TP53*-wildtype BCs, while M2 macrophages that depress inflammation and encourage tissue repair had lower proportions in *TP53*-mutated BCs. This finding indicates that *TP53* mutations promote inflammatory infiltrates and inhibit tissue repair in BC, which may contribute to higher invasion of *TP53*-mutated BCs compared to *TP53*-wildtype BCs.

**Figure 4.**
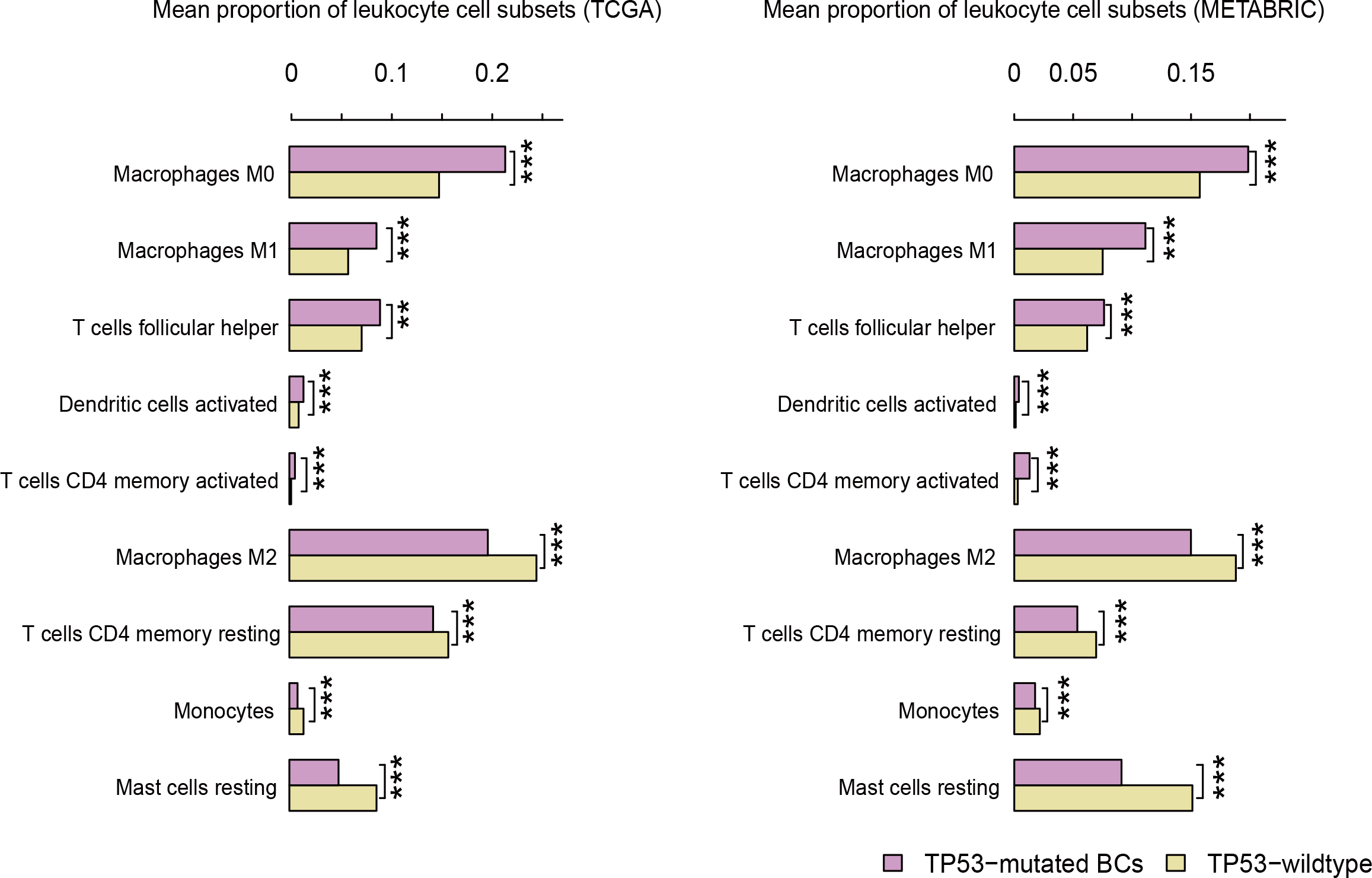
*TP53*-mutated breast cancers (BCs) have significantly different leukocyte cell subset infiltrates compared to *TP53*-wildtype BCs.

### Immune activities are associated with activities of p53-regulated pathways in BC

P53 plays important roles in regulating the cancer-associated pathways such as cell cycle, apoptosis, DNA damage repair, autophagy, metabolism, inflammation, epithelial–mesenchymal transition (EMT), angiogenesis, and metastasis [50]. Accordingly, *TP53* mutations often result in disturbances of the pathways regulated by p53 [58]. Indeed, we found that a number of p53-regulated pathways showed significantly differential activities between *TP53*-mutated and *TP53*-wildtype BCs such as the p53, cell cycle, apoptosis, Jak-STAT, NOD-like receptor, glycolysis, and Wnt pathways showing significantly higher activities in *TP53*-mutated BCs than in *TP53*-wildtype BCs (Mann-Whitney U test, P<0.05). Interestingly, most of the analyzed immune gene-sets had significantly positive correlations with the pathways upregulated in *TP53*-mutated BCs except the CT antigen gene-set with a negative correlation (Figure 5A). These results suggest that *TP53* mutations may cause disturbances of the p53-regulated pathways thereby contributing to the elevated immune activities in *TP53*-mutated BCs.

**Figure 5.**
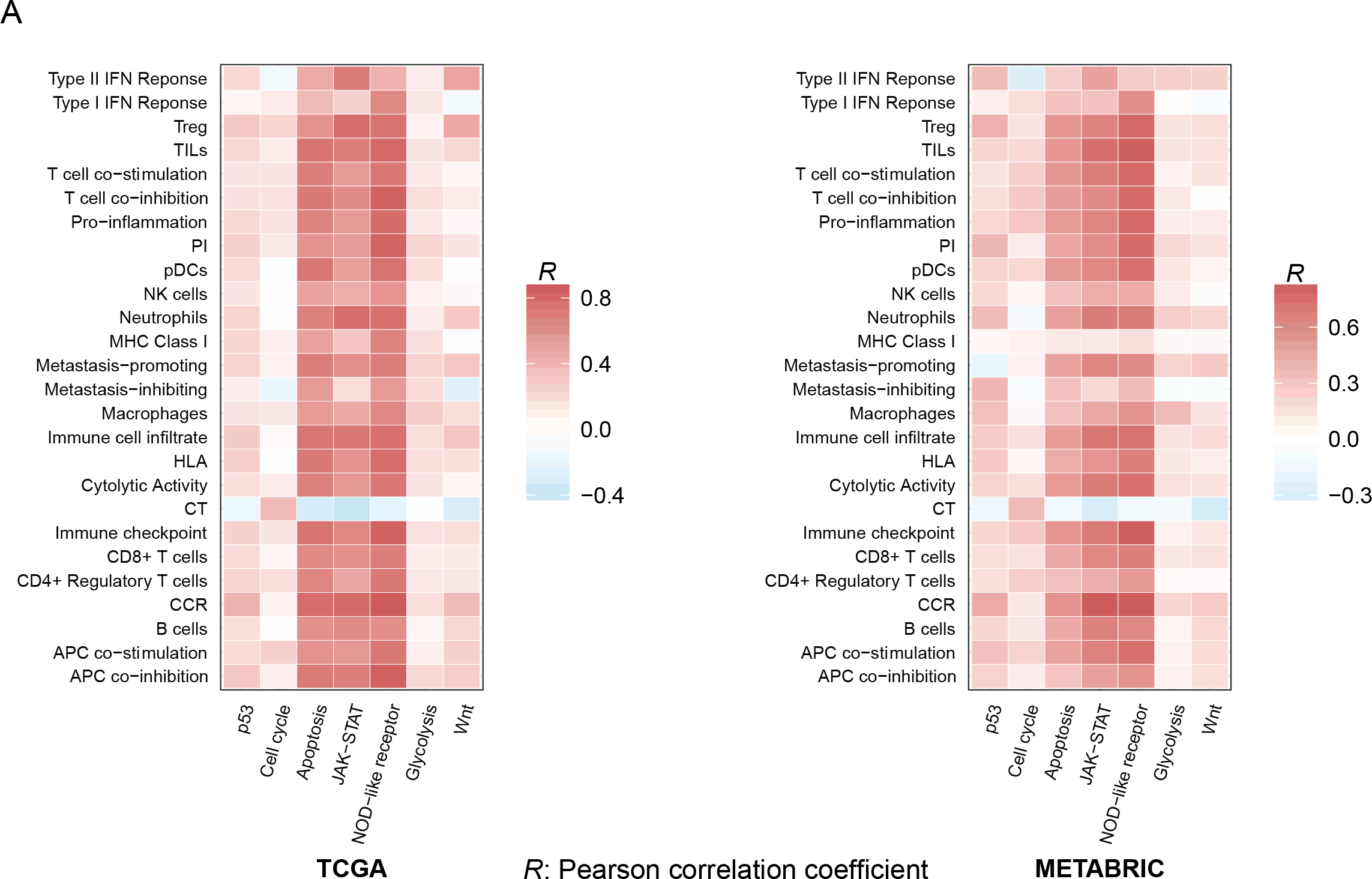
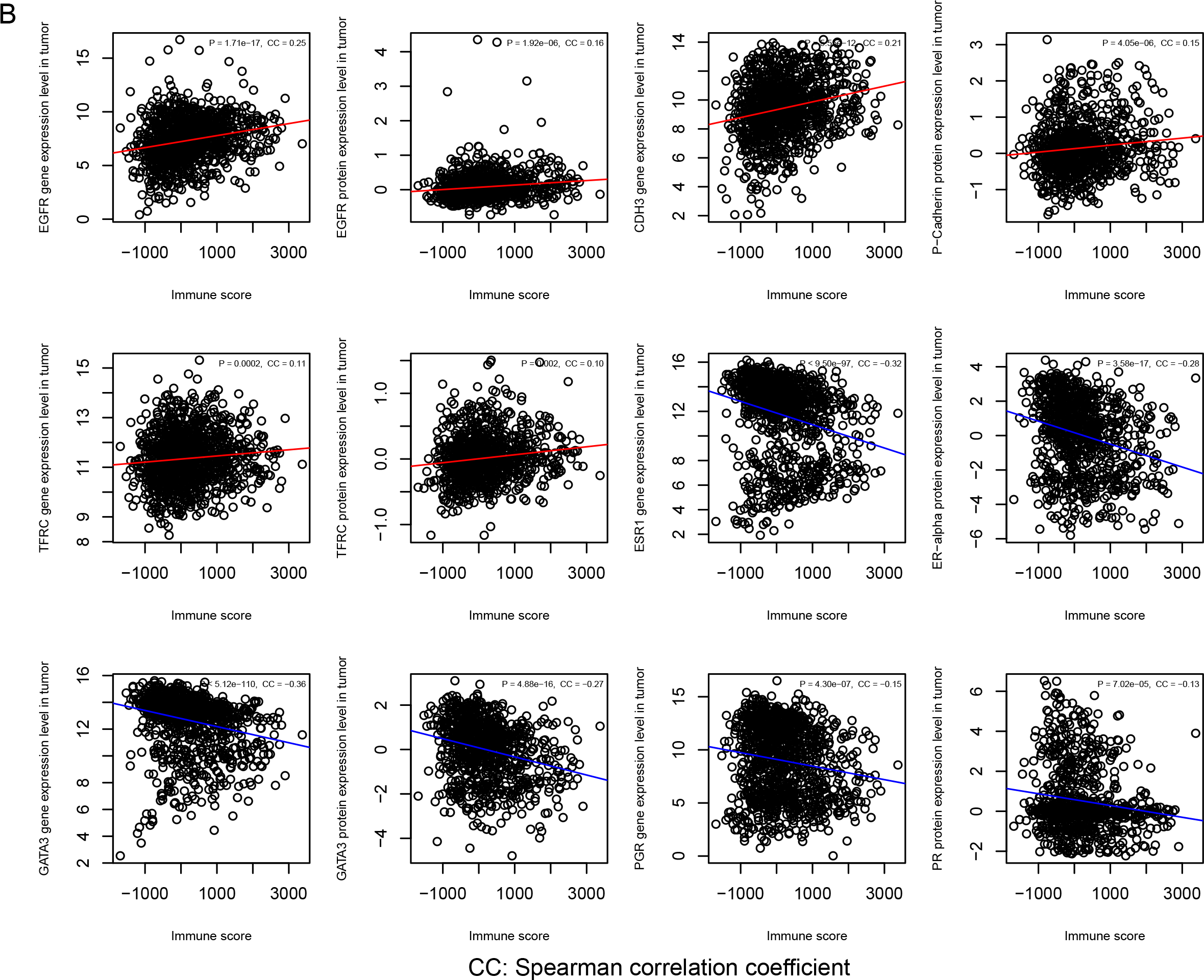
Immune activities significantly correlate with p53-regulated pathways, genes, and proteins in BC. A. Immune activities likely positively correlate with the p53-mediated pathways that show higher activities in *TP53*-mutated BCs than in *TP53*-wildtype BCs. B. Immune activities positively correlate with *EGFR, CDH3*, and *TFRC*, and their protein products that are upregulated in *TP53*-mutated BCs, while negatively correlate with *ESR1, GATA3*, and *PCR*, and their protein products that are downregulated in *TP53*-mutated BCs relative to *TP53*-wildtype BCs.

### Genes and proteins differentially expressed between *TP53*-mutated and *TP53*-wildtype BCs have significant expression correlation with immune activities in BC

Based on the gene and protein expression data in TCGA, we identified the genes and proteins that were differentially expressed between *TP53*-mutated and *TP53*-wildtype BCs (Student’s *t* test, FDR<0.05). Of these, 10 genes (*EGFR, CDH3, TFRC, CCNE1, CDK1, CDKN2A, CHEK1, FOXM1, NDRG1*, and *STMN1*) and their protein products had significantly higher expression levels in *TP53*-mutated BCs, and 8 genes (*ESR1, GATA3, PGR, AR, ERBB3, BCL2, IGF1R*, and *CCND1*) and their protein products had significantly lower expression levels in *TP53*-mutated BCs. Interestingly, almost all the immune gene-sets had significantly positive expression correlation with the 10 genes and proteins upregulated in *TP53*-mutated BCs, while significantly negative expression correlation with the 8 genes and proteins downregulated in *TP53*-mutated BCs (Spearman correlation, FDR<0.05; Additional file 7, Figures S4A, S4B). Moreover, the 10 genes and proteins likely had positive expression correlation with the immune cell infiltration scores in BC, while the 8 genes and proteins likely had negative expression correlation with the immune cell infiltration scores (Figure 5B; Additional file 7, Figure S4C). These results showed that the expression of these molecules was associated with elevated or depressed immune activities in BC.

### Association of immune activities with clinical outcomes in BC

Among the analyzed 26 immune gene-sets, 17 and 15 gene-sets showed significant correlation of enrichment levels with survival (OS and/or disease free survival (DFS)) prognosis in *TP53*-mutated and *TP53*-wildtype BCs, respectively (log-rank test, unadjusted P<0.05) (Figure 6A; Additional file 8, Figures S5A, S5B). Strikingly, elevated enrichment of the 17 gene-sets was consistently associated with better prognosis in *TP53*-mutated BCs. In contrast, of the 15 gene-sets, 10 and 5 were positively and negatively associated with prognosis in *TP53*-wildtype BCs, respectively. The B cell, cytolytic activity, T cell co-inhibition, immune checkpoint, TILs, CCR, HLA, and pro-inflammatory gene-sets showed positive correlations with prognosis consistently in the *TP53*-mutated and *TP53*-wildtype BCs. However, the CD4+ regulatory T cell gene-set showed a positive correlation with DFS prognosis in *TP53*-mutated BCs, while a negative correlation in *TP53*-wildtype BCs (Additional file 2, Table S10).

**Figure 6.**
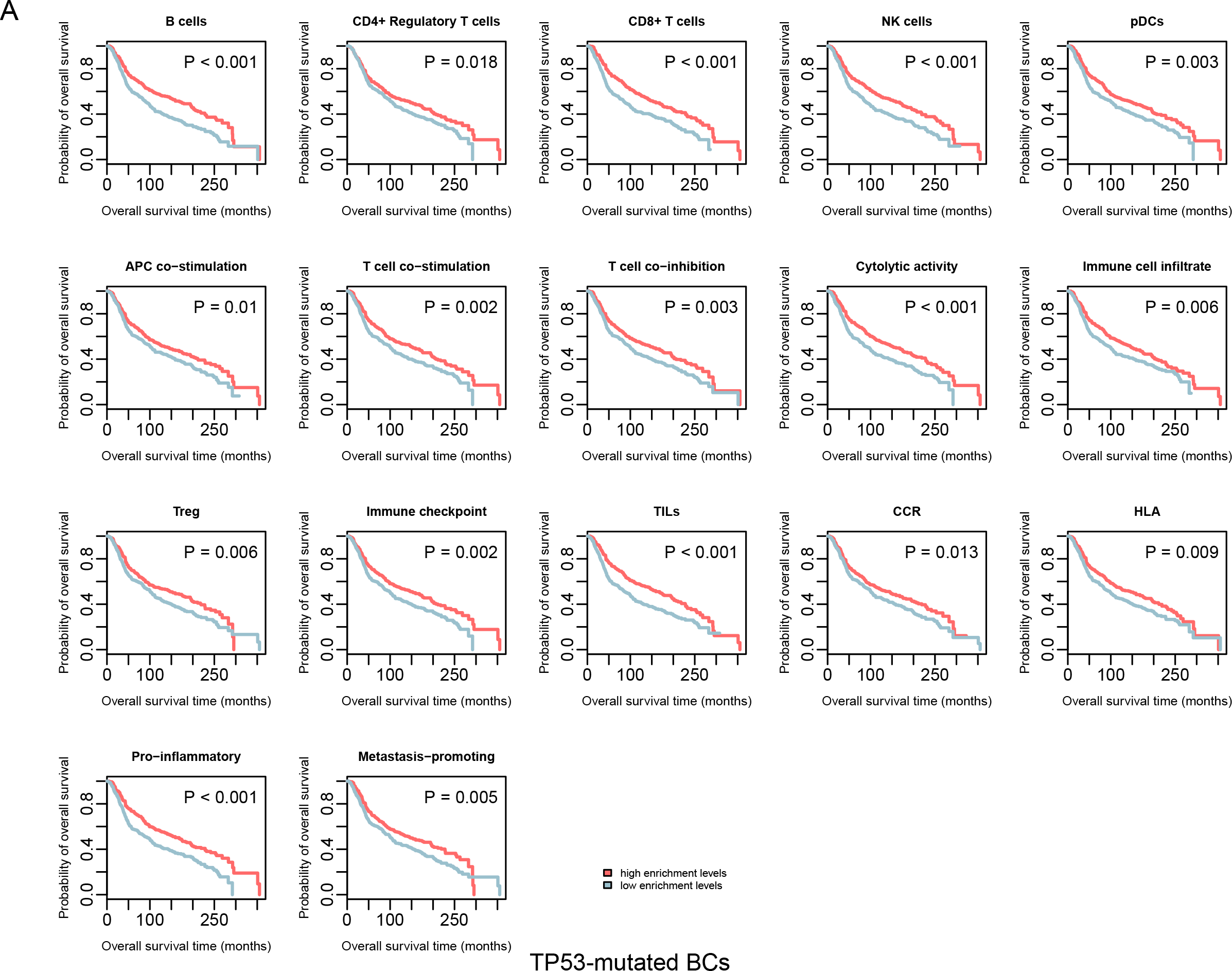
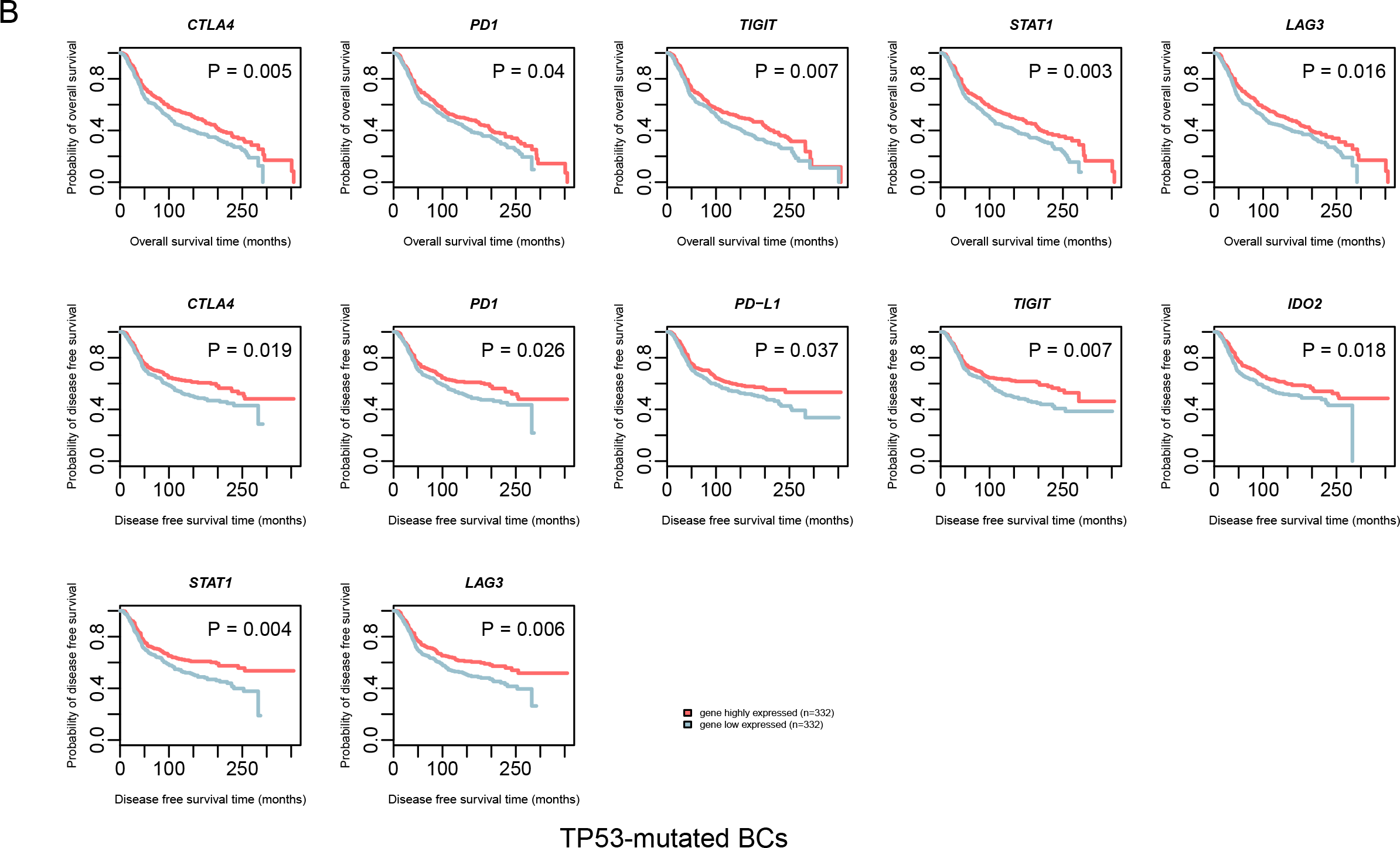
Immune activities are positively associated with survival prognosis in *TP53*-mutated BCs. A. Kaplan-Meier survival curves show that elevated enrichment of immune gene-sets is associated with better survival prognosis in *TP53*-mutated BCs. B. Kaplan-Meier survival curves show that elevated expression of immune genes is associated with better survival prognosis in *TP53*-mutated BCs. The log-rank test P<0.05 indicates the significance of survival-time differences between two classes of patients.

Furthermore, we found a substantial number of immune-related genes whose expression was associated with survival prognosis in *TP53*-mutated and/or *TP53*-wildtype BCs. For example, the immune checkpoint genes *CTLA4, PD1, PD-L1, PD-L2*, and *TIGIT*, and the CD8+ T cell marker gene *CD8A* were positively associated with prognosis in both *TP53*-mutated and *TP53*-wildtype BCs (Figure 6B; Additional file 8, Figure S5C). Besides, some genes were positively associated with prognosis exclusively in *TP53*-mutated or *TP53*-wildtype BCs such as *IL10, CD247, GZMA, GZMB, CD276, CCR4* and *CCR7* (Additional file 2, Table S11). Moreover, some genes showed inconsistent correlations with prognosis between *TP53*-mutated and *TP53*-wildtype BCs, such as *IDO2, STAT1*, and *LAG3* having positive correlations with prognosis in *TP53*-mutated BCs while negative correlations with prognosis in *TP53*-wildtype BCs (Figure 6B; Additional file 2, Table S11; Additional file 8, Figure S5C). The mechanism underlying these discrepancies may lie in that *TP53* mutations alter the TIM in BC.

### *TP53* mutations increase the expression of MHC class I genes in MCF-7 cells

We used a pair of isogenic BC cell lines with different p53 status (MCF-7 p53-wildtype and MCF-7 p53-mutant), and evaluated MHC class I gene expression levels in both cell lines. The MHC Class I genes (*HLA-A, HLA-B, HLA-C*, and *B2M*) had significantly higher mRNA expression levels in p53-mutant MCF-7 cells than in p53-wildtype MCF-7 cells, demonstrated by real-time qPCR (Figure 7A). These experimental results verified that *TP53* mutations increased the expression of HLA molecules in BC.

**Figure 7.**
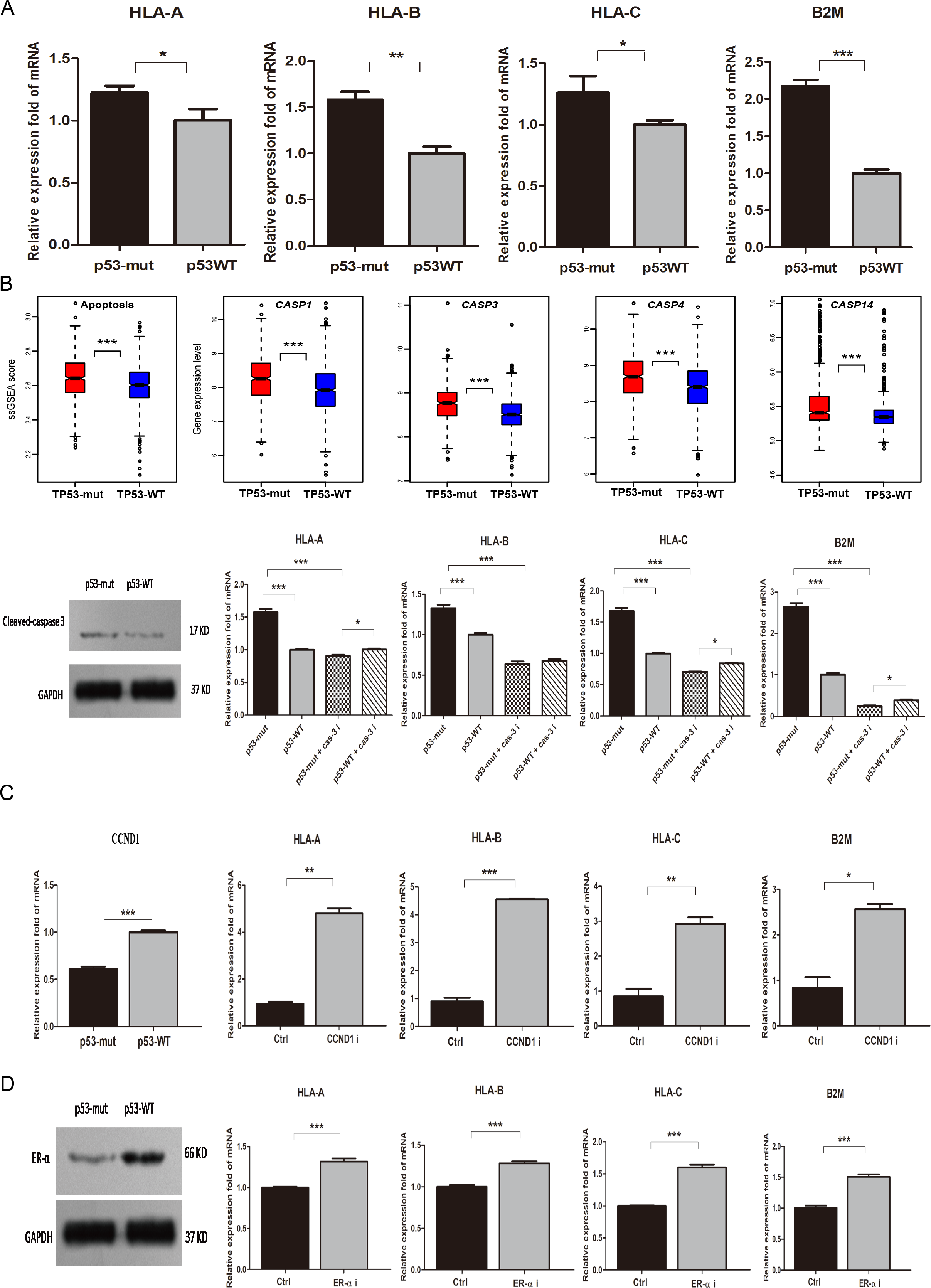
Expression of MHC Class I genes is significantly higher in p53-mutant MCF-7 cells than in p53-wildtype MCF-7 cells, and is regulated by the cell cycle, apoptosis, and estrogen receptor (ER) activities. A. MHC Class I genes (HLA-A, HLA-B, HLA-C, and B2M) have significant higher mRNA expression levels in p53-mutant MCF-7 cells than in p53-wildtype MCF-7 cells, evident by real-time quantity PCR. B. Promotion of apoptosis increases expression of MHC Class I genes, evident by both computational and experimental analyses. C. Inhibition of cell cycle increases expression of MHC Class I genes. D. Inhibition of ER alpha increases expression of MHC Class I genes. p53-WT: p53-wildtype. p53-mut: p53-mutant. CCND1 i: CCND1 inhibitor. ERα i: ER alpha inhibitor.

### *TP53* mutations increase the expression of MHC class I genes *via* regulation of apoptosis in BC

P53 plays an important role in regulation of apoptosis [59]. Surprisingly, our bioinformatics analyses showed that *TP53*-mutated BCs had significantly higher activity of the apoptosis pathway than *TP53*-wildtype BCs (Figure 7B), and *TP53*-mutated BCs more highly expressed apoptosis-inducing caspases than *TP53*-wildtype BCs such as *CASP1, CASP3, CASP4*, and *CASP14* (Figure 7B). Furthermore, our experiments verified that caspase-3 expression markedly increased in p53-mutant MCF-7 cells versus p53-wildtype MCF-7 cells (Fig 7B). We treated both p53-mutant and p53-wildtype MCF-7 cells with the caspase-3 inhibitor Z-DEVD-FMK, and found that the MHC Class I genes had markedly decreased expression in both p53-mutant and p53-wildtype MCF-7 cells (Figure 7B). Interestingly, we observed that p53-mutant MCF-7 cells more lowly expressed three of the four MHC Class I genes than p53-wildtype MCF-7 cells after they were treated with Z-DEVD-FMK (Figure 7B). These data indicate that apoptosis may have an appreciable effect on tumor immunity, and *TP53* mutations alter tumor immunogenicity *via* regulation of apoptosis.

### *TP53* mutations increase the expression of MHC class I genes *via* regulation of cell cycle in BC

Our bioinformatics analyses showed that *CCND1* (cyclin D1), a regulator of cyclin-dependent kinases, was downregulated in *TP53*-mutated BCs versus *TP53*-wildtype BCs. Furthermore, our experiments verified that *CCND1* had significantly lower mRNA expression levels in p53-mutant MCF-7 cells than in p53-wildtype MCF-7 cells (Figure 7C). We treated p53-wildtype MCF-7 cells with the cyclin D1 inhibitor abemaciclib, and observed a substantial increase in the expression of MHC Class I genes in MCF-7 cells (Figure 7C). Thus, the alteration of the p53-mediated cell cycle pathway may contribute to the differential immunogenicity between *TP53*-mutated and *TP53*-wildtype BCs.

### *TP53* mutations increase the expression of MHC class I genes *via* downregulation of estrogen receptor alpha

Our bioinformatics analyses showed that both *ESR1* and its protein product estrogen receptor alpha (ERα) were downregulated in *TP53*-mutated BCs versus *TP53*-wildtype BCs. It is consistent with previous studies showing that p53 upregulated ERα expression in BC, and *TP53* mutations downregulated ERα expression [60, 61]. Furthermore, our experiments verified that ERα was more lowly expressed in p53-mutant MCF-7 cells than in p53-wildtype MCF-7 cells (Figure 7D). We treated p53-wildtype MCF-7 cells with the ERα inhibitor MPP, and observed a marked increase in the expression of MHC Class I genes in MCF-7 cells (Figure 7D). These results indicate that the downregulation of ERα may contribute to the elevated tumor immunogenicity in *TP53*-mutated BCs.

### Mutant p53 promotes migration and proliferation of NK cells co-cultured with MCF-7 cells

We used the transwell migration and EdU proliferation assay to observe the migration and proliferation of NK92 cells co-cultured with p53-mutant and p53-wildtype MCF-7 cells for 24h, respectively. We found that the number of migrated NK92 cells co-cultured with p53-mutant MCF-7 cells far exceeded the number of migrated NK92 cells co-cultured with p53-wildtype MCF-7 cells (Figure 8A). Moreover, the NK92 cells co-cultured with p53-mutant MCF-7 cells showed significantly stronger proliferation ability compared to the NK92 cells co-cultured with p53-wildtype MCF-7 cells (Figure 8B). Furthermore, we observed that the cytokines CCL4 and CCL5 had markedly higher levels in the serum containing NK92 cells co-cultured with p53-mutant MCF-7 cells than in the serum containing NK92 cells co-cultured with p53-wildtype MCF-7 cells (Figure 8C). These observations verified our computational results that *TP53*-mutated BCs had stronger activities of immune cells including NK cells, and more highly expressed a number of CCR genes including *CCL4* and *CCL5* than *TP53*-mutated BCs. These findings are also consistent with previous studies showing that cytokines such as CCL4 could induce NK cells migration [62] and that activated NK cells could secrete cytokines to mediate immune response [63].

**Figure 8.**
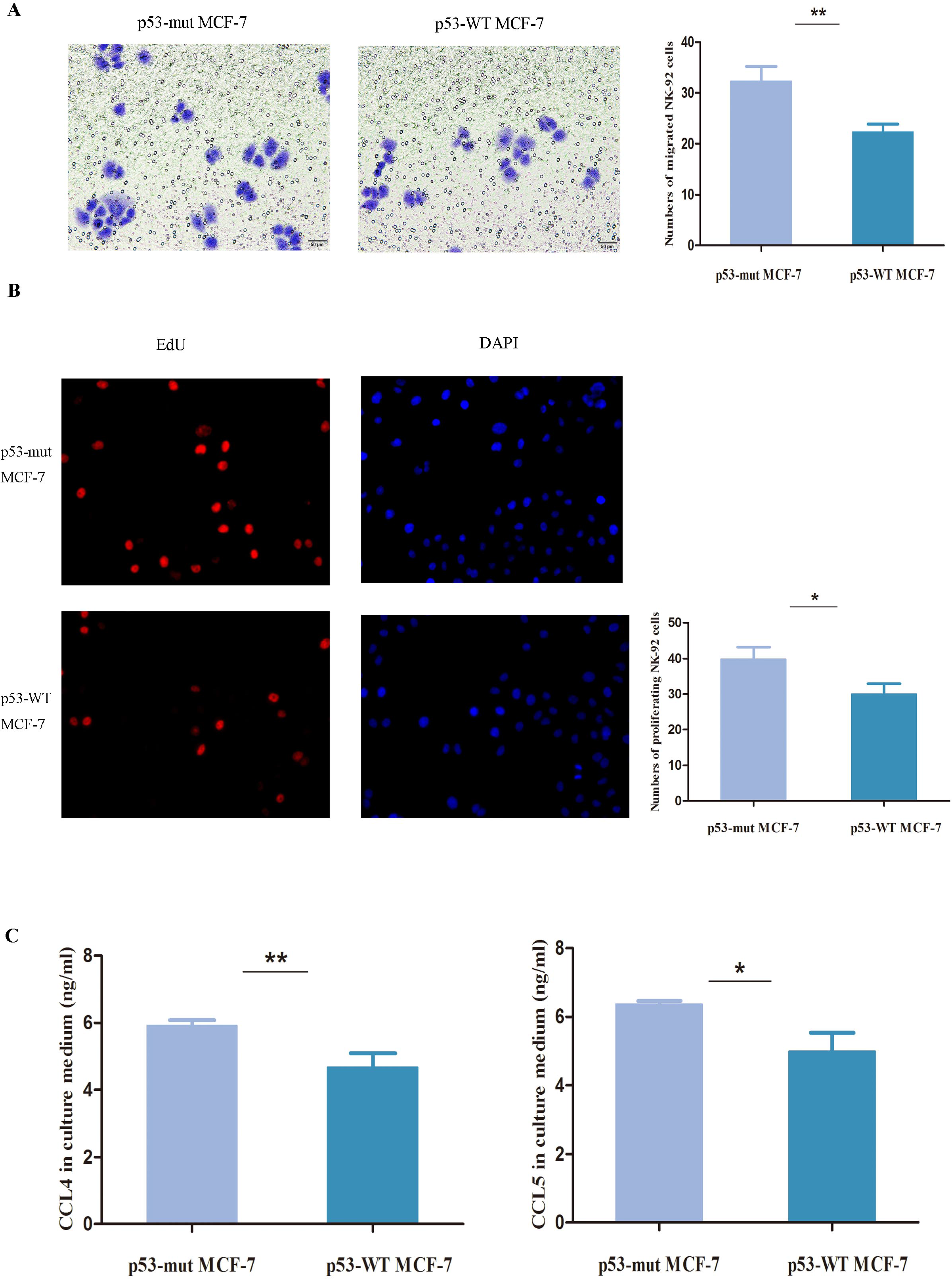
Mutant p53 promotes migration and proliferation of NK cells co-cultured with MCF-7 cells. A. NK92 cells co-cultured with p53-mutant MCF-7 cells show stronger migration ability than NK92 cells co-cultured with p53-wildtype MCF-7 cells, evident by transwell migration assay. B. NK92 cells co-cultured with p53-mutant MCF-7 cells show stronger proliferation ability than NK92 cells co-cultured with p53-wildtype MCF-7 cells, evident by EdU proliferation assay. C. The cytokines CCL4 and CCL5 have markedly higher levels in the serum involving NK92 cells co-cultured with p53-mutant MCF-7 cells than in the serum involving NK92 cells co-cultured with p53-wildtype MCF-7 cells, evident by quantitative enzyme-linked immunosorbent assay (ELISA).

## Discussion

In the present study, we performed a comprehensive portrait of the associations between *TP53* mutations and immune activities in BC. We found that *TP53*-mutated BCs showed significantly higher activities of a wide variety of immune cells, functions, and pathways than *TP53*-wildtype BCs (Figure 1A). Moreover, *TP53*-mutated BCs had significantly higher levels of immune cell infiltration than *TP53*-wildtype BCs in terms of immune scores and clinical pathological data. Furthermore, we found that *TP53*-mutated BCs had higher proportions of activated immune cell subsets and lower proportions of resting immune cell subsets compared to *TP53*-wildtype BCs within the TME. These findings consistently demonstrate that *TP53* mutations are associated with stronger immune activities in BC.

*TP53*-mutated BCs showed significant differences in clinical features compared to *TP53*-wildtype BCs. Typically, *TP53*-mutated BCs contained a higher proportion of ER-, PR-, HER2+, or triple-negative/basal-like BCs compared to *TP53*-wildtype BCs (Figure 1A). Previous studies have showed that the ER-and HER2+ features were associated stronger immunogenic activity in BC [64]. Thus, both features may contribute to the higher immune activity of *TP53*-mutated BCs as compared to *TP53*-wildtype BCs. However, when we compared enrichment levels of the 26 immune gene-sets between *TP53*-mutated and *TP53*-wildtype BCs within the ER+ subtype of BC, we found that almost all these gene-sets had significantly higher enrichment levels in *TP53*-mutated ER+ BCs than in *TP53*-wildtype ER+ BCs (Additional file 3, Table S12). Similarly, *TP53*-mutated HER2- BCs had significantly higher enrichment levels of almost all these immune gene-sets than *TP53*-wildtype HER2- BCs (Additional file 3, Table S13). These results indicate that the *TP53* mutation itself is capable of contributing to the distinct immunogenic activity between *TP53*-mutated and *TP53*-wildtype BCs. In fact, our experimental results verified that mutant p53 could increase immunogenic activity in BC. Furthermore, our computational and experimental results suggest that *TP53* mutations may alter immunogenic activity in BC *via* regulation of the p53-mediated pathways such as cell cycle, apoptosis, Wnt, Jak-STAT, NOD-like receptor, and glycolysis. It should be noted that our findings appear to be contrary to previous studies showing that p53 increased immunogenic activity in some cancers such as colon cancer [49] and lymphom [65]. However, a number of studies have showed that p53 functions in a context-dependent fashion [50, 51, 65]. Thus, these distinct effects of p53 in regulating tumor immunity could be attributed to the different cellular contexts.

One tumor sample may involve a certain percent of non-tumor cells such as normal cells and stromal cells. To exclude the impact of non-tumor cells on tumor immunity, we selected the BC samples composed of 100% tumor cells based on the TCGA BC pathological slides data. We found that almost all the 26 immune gene-sets showed significantly higher enrichment levels in the *TP53*-mutated class of these samples than in the *TP53*-wildtype class of these samples (Mann-Whitney U test, FDR<0.05) (Additional file 3, Table S14). Thus, the distinction of immunogenic activity between *TP53*-mutated and *TP53*-wildtype BCs indeed referred to the difference in tumor immunity.

The elevated expression of immunosuppressive, pro-inflammatory, and metastasis-promoting genes in *TP53*-mutated BCs may promote tumor invasion and lead to a worse prognosis in BC. Indeed, previous studies have shown that p53 mutations were associated with unfavorable clinical outcomes in BC [66, 67]. The METABRIC data also showed that *TP53*-mutated BCs had worse OS and DFS prognosis compared to *TP53*-wildtype BCs (Figure 9A). Moreover, *TP53*-mutated BCs more highly expressed Ki67 (a marker for cell proliferation) than *TP53*-wildtype BCs (Figure 1A), again indicating the higher aggressiveness of *TP53*-mutated BCs compared to *TP53*-wildtype BCs.

**Figure 9.**
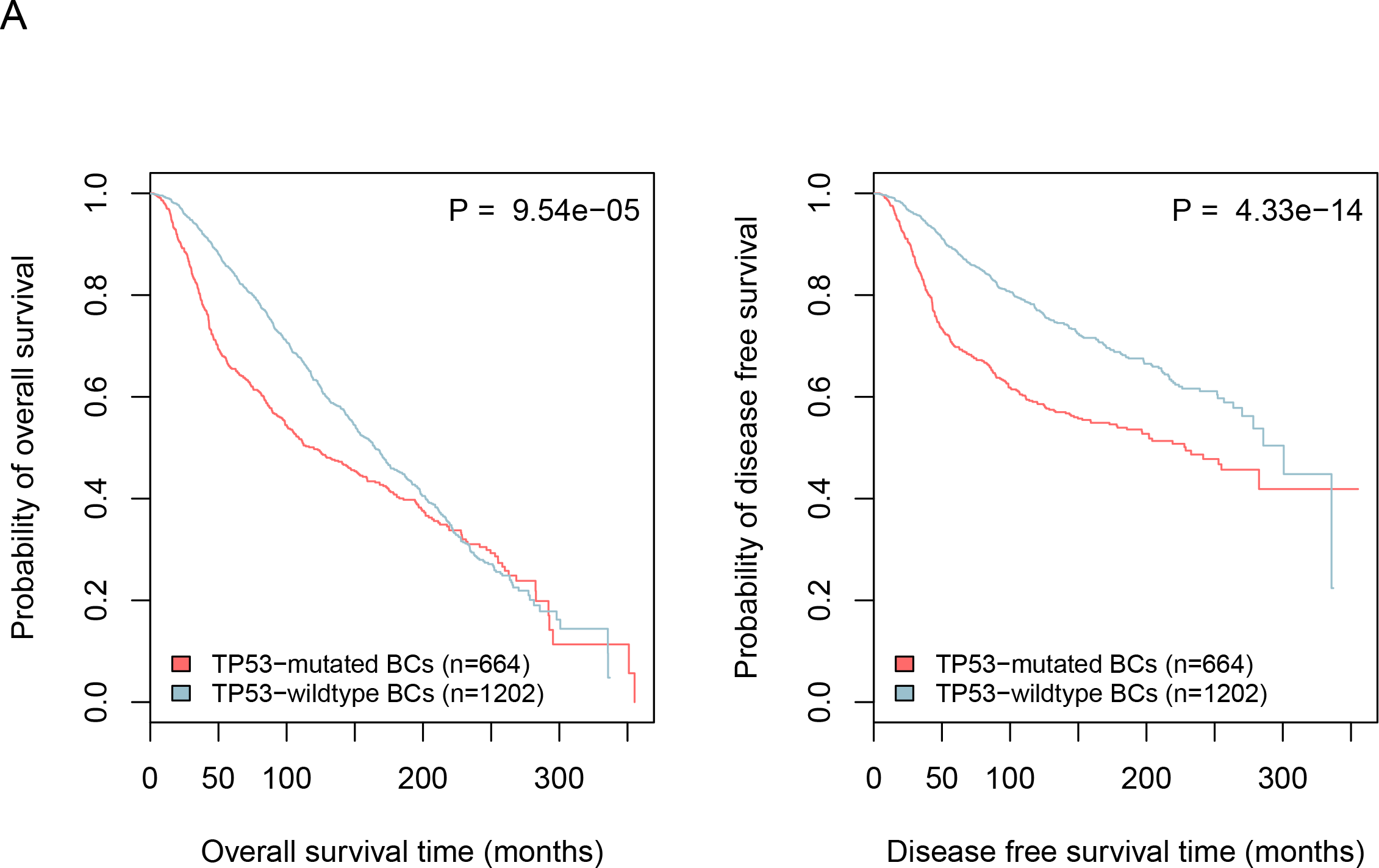
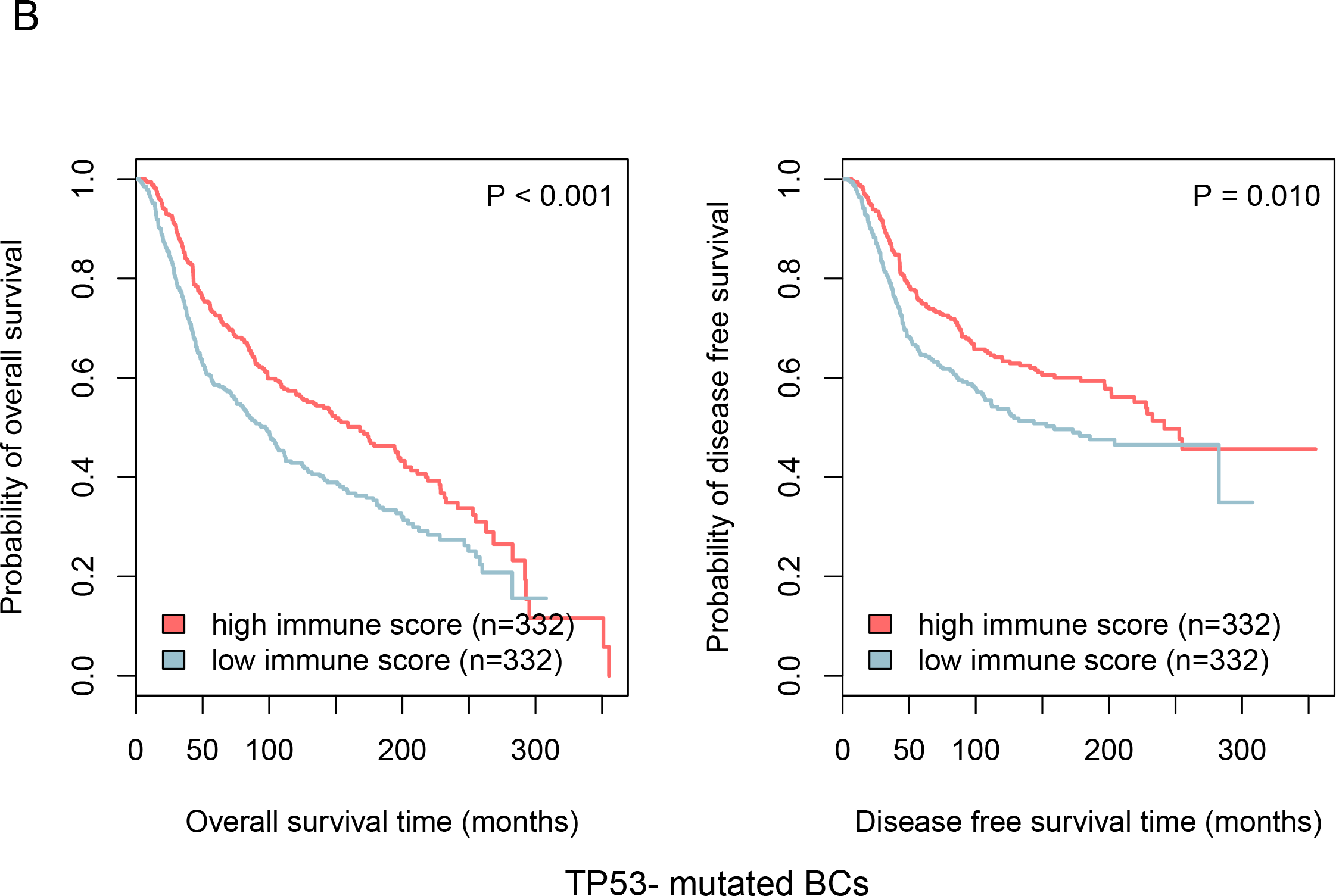
*TP53* mutations are associated unfavorable survival prognosis in BC, while higher degree of immune cell infiltration is associated with better survival prognosis in BC. A. Kaplan-Meier survival curves show that *TP53*-mutated BCs have worse survival prognosis compared to *TP53*-wildtype BCs. B. Kaplan-Meier survival curves show that higher degree of immune cell infiltration is associated with better survival prognosis in *TP53*-mutated BCs. The log-rank test P<0.05 indicates the significance of survival-time differences between two classes of patients.

Interestingly, the activities of different immune cell types, functions and pathways, and the immune cell infiltration degree were consistently positively associated with survival prognosis in *TP53*-mutated BCs (Figures 6A, 6B, 9B). It is sensible that the elevated enrichment of CD8+ T cell, B cell, NK cell, cytolytic activity, HLA, immune cell infiltrate, TILs, and CCR is associated with favorable clinical outcomes in cancer since these immune activities promote anticancer immunity. Furthermore, the observation that the elevated enrichment of Treg, immune checkpoint, pro-inflammatory, and metastasis-promoting immune activities was associated with better survival prognosis in *TP53*-mutated BCs may be due to that the elevated immunosuppressive activities are likely to promote chemotherapy sensitivity of *TP53*-mutated BCs [68]. Thus, to achieve successes in immunotherapy of *TP53*-mutated BCs, the effective combination of chemotherapy and immunotherapy may represent a promising direction [69].

Interestingly, compared to *TP53*-wildtype BCs, *TP53*-mutated BCs more highly expressed a majority of the gene targets for immunotherapy agents that are currently used in the clinic, clinical trials or preclinical development [70] (Additional file 3, Table S15). It indicates that these immunotherapy agents may be more effective against *TP53*-mutated BCs than *TP53*-wildtype BCs. In fact, several clinical trials [21, 22] have shown that immune checkpoint blockade was effective against TNBC, a BC subtype with a high *TP53* mutation rate (80%).

## Conclusions

*TP53* mutations promote immunogenic activity in breast cancer. This finding suggests that the *TP53* mutation status could be a useful biomarker for stratifying BC patients responsive to immunotherapy.

## List of abbreviations

BC: Breast Cancer
BRCA: BReast invasive CArcinoma
CCR: Cytokine and Cytokine Receptor
CT: Cancer Testis
DFS: Disease Free Survival
FDR: False Discovery Rate
GSVA: gene-set variation analysis
HLA: Human Leukocyte Antigen
iDCs: immature dendritic cells
ME: mean expression
MHC: Major Histocompatibility Complex
MMR: mismatch-repair
OR: Odds Ratio
OS: Overall Survival
pDCs: plasmacytoid Dendritic Cells
PI: Parainflammation
ssGSEA: single-sample gene-set enrichment analysis (ssGSEA)
TCGA: The Cancer Genome Atlas
Tfh: T Follicular Helper
TILs: Tumor-Infiltrating Lymphocytes
TIM: tumor immune microenvironment
TMB: Tumor Mutation Burden
TME: tumor immune microenvironment
TNBC: Triple-Negative Breast Cancer
Treg: Regulatory T

## Declarations

### Ethics approval and consent to participate

Ethical approval was waived since we used only publicly available data and materials in this study.

### Consent for publication

Not applicable.

### Availability of data and material

We downloaded TCGA RNA-Seq gene expression profiles (Level 3), gene somatic mutations (Level 3), SCNAs (Level 3), protein expression profiles (Level 3), and clinical data for BC from the genomic data commons data portal (https://portal.gdc.cancer.gov/), and the METABRIC gene expression profiles, gene somatic mutations, SCNAs, and clinical data from cBioPortal (http://www.cbioportal.org). We performed all the computational and statistical analyses using R programming (https://www.r-project.org/).

## Authors’ contributions

ZL performed data analyses and experiments, and wrote the manuscript. ZJ performed data analyses and helped prepare for the manuscript. YG performed experiments. LW performed experiments. CC performed data analyses. XW conceived the research, designed analysis strategies, performed data analyses, and wrote the manuscript. All the authors read and approved the final manuscript.

## Acknowledgements

We thank Dr. Jiaqi Ge for his valuable suggestions on our experimental design.

## Additional files

### Additional file 1: Tables S1-9

Table S1. Comparison of expression levels of the immune cell types and functional marker genes and gene-sets between two classes of samples.

Table S2. Comparison of expression levels of tumor-infiltrating lymphocytes genes and gene-set between two classes of samples.

Table S3. Comparison of expression levels of the Treg genes and gene-set between two classes of samples.

Table S4. Comparison of expression levels of the immune checkpoint genes and gene-set between two classes of samples.

Table S5. Comparison of expression levels of the cytokine and cytokine receptor genes and gene-set between two classes of samples

Table S6. Comparison of expression levels of the cancer-testis genes and gene-set between two classes of samples.

Table S7. Comparison of expression levels of the HLA genes and gene-set between two classes of samples.

Table S8. Comparison of expression levels of the inflammation-promoting and parainflammation (PI) genes and gene-sets between two classes of samples.

Table S9. Comparison of expression levels of the metastasis-promoting and metastasis-inhibiting genes and gene-sets between two classes of samples.

### Additional file 2: Tables S10-11

Table S10. Comparisons of ssGSEA scores of immune gene-sets between *TP53*-mutated and *TP53*-wildtype BCs, and their associations with survival prognosis in BC.

Table S11. Comparisons of expression levels of immune genes between *TP53*-mutated and *TP53*-wildtype BCs, and their associations with survival prognosis in BC.

### Additional file 3: Tables S12-16

Table S12. Comparisons of enrichment levels of immune gene-sets between *TP53*-mutated and *TP53*-wildtyped BCs within the ER+ subtype.

Table S13. Comparisons of enrichment levels of immune gene-sets between *TP53*-mutated and *TP53*-wildtyped BCs within the HER2- subtype.

Table S14. Comparisons of enrichment levels of immune gene-sets between *TP53*-mutated and *TP53*-wildtyped BCs within the 100% tumor purity.

Table S15. Comparisons of expression levels of the genes targeted by immunotherapy agents in clinical use or trials or in preclinical development between *TP53*-mutated and *TP53*-wildtyped BCs.

Table S16. Primer sequences used for real time quantity PCR.

### Additional file 4

Figure S1. *TP53*-mutated breast cancers (BCs) have increased immune activities compared to *TP53*-wildtype BCs. A. Heatmap showing the ssGSEA scores of 26 immune gene-sets in *TP53*-mutated and *TP53*-wildtype BCs (TCGA). ssGSEA: single-sample gene-set enrichment analysis. TNBC: triple negative breast cancer. Red color indicates higher enrichment levels (ssGSEA scores) of gene-sets, and blue color indicates lower enrichment levels of gene-sets in the heatmap. *RB1* are more frequently mutated in *TP53*-mutated BCs while *CDH1, GATA3, MAP3K1*, and *PIK3CA* are more frequently mutated in *TP53*-wildtype BCs (Fisher’s exact test, P<0.05). The black vertical lines in the bars beside gene symbols indicate that the genes are mutated in corresponding samples. The black vertical lines in the bar beside “TNBC” indicate that the sample is a TNBC. The black vertical lines in the bars beside “ER”, “PR”, and “HER2” indicate that the sample is ER-, PR-, or HER2-. B. Heatmap showing *TP53*-mutated BCs likely more highly express immune cell types and functional marker genes than *TP53*-wildtype BCs (TCGA). NK: natural killer. pDCs: plasmacytoid dendritic cells. MHC: histocompatibility complex. APC: antigen-presenting cell. C. Most of the immune cell types and functional marker gene-sets show higher enrichment levels in *TP53*-mutated BCs than in *TP53*-wildtype BCs. D. Heatmap showing *TP53*-mutated BCs likely more highly express tumor-infiltrating lymphocytes (TILs) genes than *TP53*-wildtype BCs. E. *TP53*-mutated BCs have higher enrichment levels of the TILs gene-set than *TP53*-wildtype BCs. F. Heatmap showing *TP53*-mutated BCs likely more highly express Treg and immune checkpoint genes than *TP53*-wildtype BCs (TCGA). G. *TP53*-mutated BCs have higher enrichment levels of the Treg and immune checkpoint gene-sets than *TP53*-wildtype BCs. H. A number of important immune checkpoint genes are more highly expressed in *TP53*-mutated BCs than in *TP53*-wildtype BCs. Treg: regulatory T cell. Red color indicates higher gene expression levels, and blue color indicates lower gene expression levels in the heatmaps (it applies to all the other heatmaps). *: P < 0.05; **: P < 0.01; ***: P < 0.001, and it applies to all the other figures.

### Additional file 5

Figure S2. *TP53*-mutated breast cancers (BCs) have higher immune activities of cytokine and cancer-testis (CT) antigens than *TP53*-wildtype BCs. A. Heatmaps showing *TP53*-mutated BCs likely more highly express cytokine and cytokine receptor (CCR) genes than *TP53*-wildtype BCs. B. *TP53*-mutated BCs have higher enrichment levels of the CCR gene-set than *TP53*-wildtype BCs. C. A number of CT antigen genes are upregulated in *TP53*-mutated BCs relative to *TP53*-wildtype BCs and normal tissue, and encode the CT antigens that are potential targets for developing cancer vaccines. D. *TP53*-mutated BCs have higher enrichment levels of the CT antigen gene-set than *TP53*-wildtype BCs and normal tissue.

### Additional file 6

Figure S3. *TP53*-mutated breast cancers (BCs) have higher activities of HLA, pro-inflammation, and parainflammation (PI) than *TP53*-wildtype BCs. A. Heatmaps showing *TP53*-mutated BCs likely more highly express HLA genes than *TP53*-wildtype BCs in TCGA. B. Heatmaps showing *TP53*-mutated BCs likely more highly express pro-inflammatory and PI genes than *TP53*-wildtype BCs. C. Most of the pro-inflammatory genes are upregulated in *TP53*-mutated BCs relative to *TP53*-wildtype BCs.

### Additional file 7

Figure S4. The genes and their protein products have significant expression correlation with immune activities in BC that are differentially expressed between *TP53*-mutated and *TP53*-wildtype breast cancers (BCs). A. The genes *TFRC, CDH3*, and *EGFR* show positive expression correlation with immune activities in BC. B. The genes *PGR, ESR1, ERBB3, BCL2*, and *AR* show negative expression correlation with immune activities in BC. C. *AR, ERBB3, BCL2*, and *IGF1R*, and their protein products consistently show negative expression correlation with immune scores in BC.

### Additional file 8

Figure S5. Association of immune activities with clinical outcomes in breast cancers (BCs). A. Kaplan-Meier survival curves show that elevated enrichment of immune gene-sets is associated with better survival prognosis in *TP53*-mutated BCs. B. Kaplan-Meier survival curves show that elevated enrichment of immune gene-sets is associated with better or worse survival prognosis in *TP53*-wildtype BCs. C. The genes *IDO2, STAT1*, and *LAG3* have negative expression correlation with survival prognosis in *TP53*-wildtype BCs.

